# Integrating bacterial genomics and host transcriptomics links *Burkholderia pseudomallei* variation to melioidosis mortality and neutrophil responses

**DOI:** 10.64898/2026.11.06.764829

**Authors:** Arin Wongprommoon, Phumrapee Boonklang, Katie L. Burnham, Chalita Chomkatekaew, Kesorn Angchagun, Gumphol Wongsuvan, Thatsanun Ngernseng, Direk Limmathurotsakul, Tanupat Boonchalermvichien, Taylor D. Coston, Rungnapa Phunpang, Thatcha Yimthin, Patpong Rongkard, Wanseon Lee, Barbara Kronsteiner, Julian Knight, Jukka Corander, Sharon J. Peacock, Julian Parkhill, Nicholas P. J. Day, Susanna J. Dunachie, T. Eoin West, Narisara Chantratita, Nicholas R. Thomson, Emma E. Davenport, Claire Chewapreecha

## Abstract

Bacterial genetic variation contributes to the outcome of infection, but its relationship with host immune responses is rarely investigated in the same patients, limiting the biological interpretation of pathogen-associated disease severity. Here, we integrated pathogen genomics, clinical outcomes and host transcriptomics to investigate host-pathogen interactions in melioidosis, a frequently fatal infection caused by *Burkholderia pseudomallei*. We analysed prospective clinical data from 2,142 patients, whole-genome sequences from 2,101 infecting isolates and 646 paired host blood transcriptomes across five cohorts collected between 2012 and 2023. Infections were typically dominated by a single strain, with limited within-host diversification, supporting the use of an infecting isolate genome to investigate pathogen- associated clinical outcomes. Bacterial genome-wide association studies identified one locus associated with binary mortality at genome-wide significance. Incorporating patient age, sex, body mass index and Charlson Comorbidity Index increased this to four, while modelling time to death as a continuous phenotype identified seven loci. Integration with paired host transcriptomes linked mortality-associated bacterial genetic variation to imputed signatures of increased immature neutrophils and reduced neutrophil degranulation. An intergenic region upstream of the polyketide synthesis locus *BPSS1006*-*BPSS1011* was additionally associated with downregulation of host genes involved in myeloid-cell signalling and cell proliferation pathways. These findings connect naturally occurring *B. pseudomallei* genomic variation associated with mortality to distinct host immune signatures, providing insight into pathogen- host interactions underlying severe melioidosis and a framework for investigating the contribution of pathogen genetic diversity to clinical outcomes.

## Introduction

The outcome of a bacterial infection is determined by complex interactions between pathogen virulence factors and host factors that contribute to disease severity. While host characteristics, including age, comorbidities and immune status, are major determinants of disease severity^1,2^; bacterial genetic variation influences the pathogen’s capacity to invade, evade immune responses and disseminate within the host^3,4^. Bacterial genome-wide association studies (GWAS) have emerged as a powerful approach for linking genomic variation to microbial phenotypes^5^, revealing genetic determinants of antimicrobial resistance^6–8^, host carriage^9^, host adaptation^10^, and tissue tropism^11^. However, identifying bacterial genetic associations with clinical outcomes has proved more challenging. Unlike classical bacterial traits, outcomes such as mortality reflect the combined contributions of the pathogen and the infected individual. Consequently, pathogen-associated genetic effects are often small^12^ relative to variation arising from host factors; the cohort sizes achievable for individual infections can further limit statistical power^12,13^. Accounting for host characteristics as covariates in bacterial GWAS provides an established way to address this heterogeneity and has been shown to recover associations that may be missed by pathogen-only models^14^. However, statistical adjustment alone does not establish how bacterial variation relates to the host response during infection. This requires pathogen genomic data to be considered alongside molecular measurements of the host in the same patients.

Melioidosis, caused by the environmental Gram-negative bacterium *Burkholderia pseudomallei*, provides an important setting in which to investigate these relationships. The disease is endemic throughout tropical regions and remains associated with high mortality despite appropriate antimicrobial therapy^15,16^. Clinical manifestations range from localised infection to fulminant septic shock, with outcome strongly influenced by host factors including diabetes, chronic renal disease, chronic liver disease, and route of infection^17,18^. These host-associated effects account for a substantial proportion of the observed variation in mortality and may obscure bacterial genetic contributions if not explicitly modelled. At the same time, *B. pseudomallei* possesses large genomic diversity^19,20^ and encodes a broad repertoire of virulence determinants implicated in host attachment^21–23^, including pili^24^; immune evasion^24,25^; host cell invasion; and intracellular survival and dissemination^26,27^, including effectors produced by polyketide synthases^27^. Experimental studies have established roles for many of these factors in pathogenesis, but have largely focused on candidate genes, individual mechanisms, or relatively small numbers of strains^24,25,27^. Consequently, the contribution of naturally occurring bacterial genetic variation to mortality has not been systematically investigated at genome scale. Moreover, whether outcome-associated bacterial variation is also associated with differences in the human immune response during acute infection remains poorly understood.

To address these questions, we assembled prospective clinical data from 2,142 patients with culture-confirmed melioidosis, whole genome sequences from 2,101 infecting isolates and 646 paired host blood transcriptomes across five cohorts recruited in northeast Thailand between 2012 and 2023. We characterised within-host diversity to assess the representativeness of individual infecting isolates and conducted bacterial GWAS accounting for host comorbidities to identify genetic associations with mortality using both binary and time-to-event outcomes. By integrating mortality-associated bacterial genetic variation with paired host transcriptomes, we further investigated the relationship between pathogen genotype and host immune responses during acute infection. Our findings identify pathogen genotype and host immune responses during acute infection. Our findings identify bacterial genetic determinants associated with fatal melioidosis and link pathogen genetic variation to differences in host immune and cellular responses, providing a framework for investigating the contribution of pathogen diversity to clinical outcomes through its relationship with host biology.

## Results

### A causal modelling framework identifies host factors to be adjusted in mortality association studies

To define host factors that should be accounted for when identifying bacterial genetic determinants of outcome, we assembled clinical data from 2,142 patients with culture-confirmed *B. pseudomallei* infection across ten hospitals in northeast Thailand between 2012-2023 (**Figures 1a-b, Supplementary Table 1**). The cohort combined previously published datasets^28–30^ with newly recruited patients^31^, providing a large, deeply phenotyped population spanning the major referral network for melioidosis in the region. The clinical characteristics were consistent with the established epidemiology of melioidosis (**Table 1**). The median age was 54.8 years (IQR 46.0-63.5), 70.8% were male consistent with increased occupational exposure to contaminated soil and surface water^17,32^, and diabetes was the most common comorbidity (66.9%). Median body mass index (BMI) was 22.0 kg m^-2^ (IQR 19.5-24.2), slightly lower than population-level estimates for northeast Thailand^33–35^. Overall, 28-day mortality was 25.7%.

**Figure 1:**
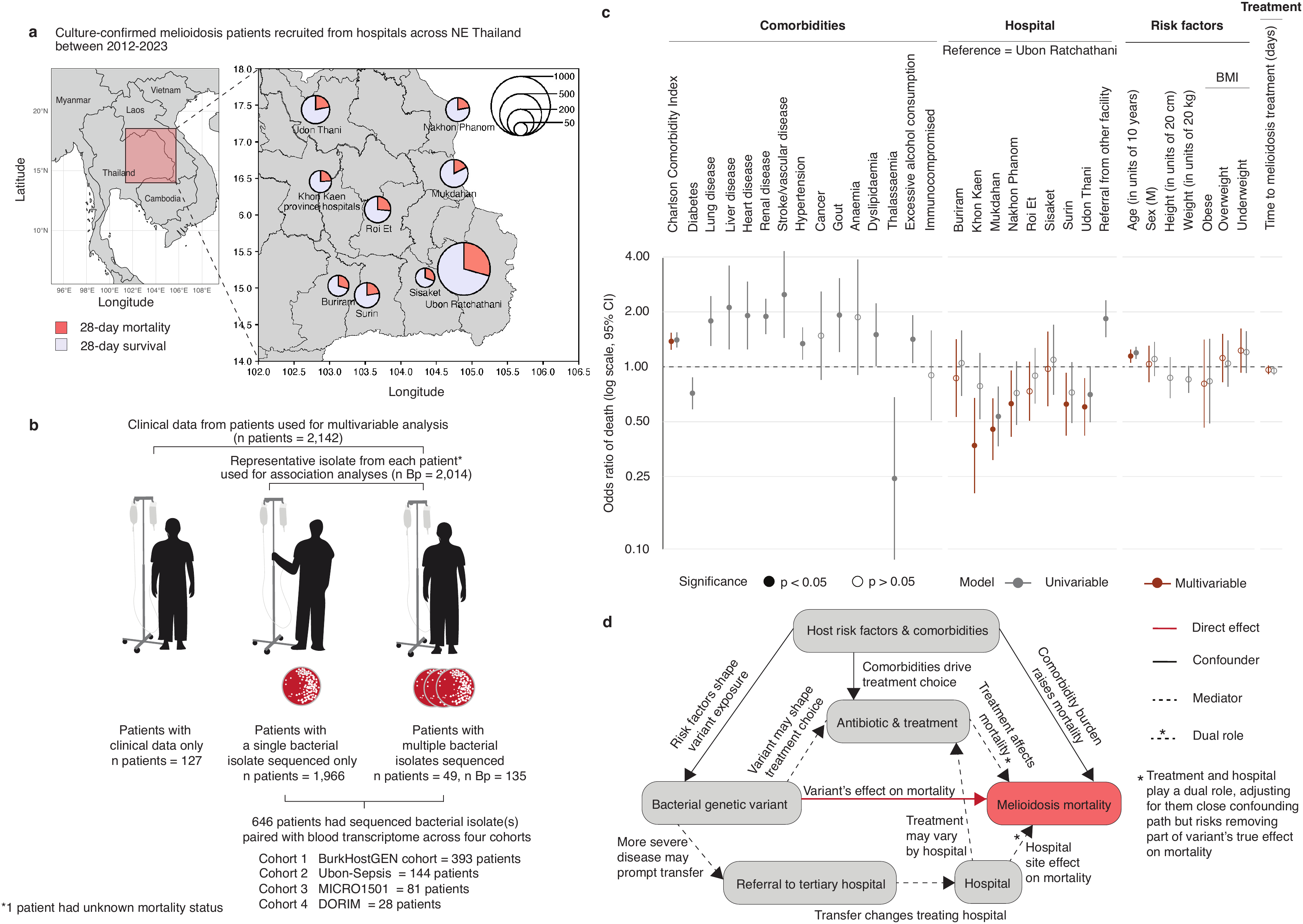
Study population, analytical framework and clinical determinants of melioidosis mortality. **(a)** Geographical distribution of culture-confirmed melioidosis patients recruited from hospitals across northeast Thailand between 2012 and 2023. The inset shows the study region within mainland Southeast Asia. Pie is proportional to the number of patients recruited at that site with pie slices showing the proportion who died within 28 days (red) or survived (lilac). **(b)** Derivation of the analysis set. **(c)** Association of host, clinical and treatment variables with 28- day mortality, from univariable (grey) and multivariable (red) logistic regression models fitted to the 2,142 patients with clinical data. Points give the odds ratio of death and bars the 95% confidence interval, on a log scale. Filled points indicate p < 0.05 and open points p > 0.05. **(d)** Directed acyclic graph setting out the assumed causal relationships between bacterial genotype, host and clinical factors, and host mortality, and used to select covariates for the adjusted models. The red edge is the effect of interest: the effect of bacterial genetic variation on mortality. Solid edges denote confounding paths, dashed edges mediating paths, and dashed edges with asterisks variables acting in both roles. Treatment and hospital are dual-role variables: each lies on a confounding path that adjustment should close and on a mediating path through which part of the true effect of the bacterial variant may travel, so adjusting for them risks removing a component of the effect being estimated.

**Table 1:** Baseline demographics of combined cohort

| **Patient characteristics** | **All (n = 2,142)** |
|---|---|
| Age in years - median (IQR) | 54.8 (46.0-63.5) |
| Age in years, groups - n (%) |  |
| < 18 | 7 (< 1%) |
| 18-30 | 74 (3.5%) |
| 31-50 | 726 (33.9%) |
| 51-70 | 1,114 (52.0%) |
| ≥ 71 | 221 (10.3%) |
| Male - n (%) | 1517 (70.8%) |
| BMI - median (IQR) | 22.0 (19.5-24.2) |
| BMI, grouped - n (%) |  |
| < 18.5 (underweight) | 347 (16.2%) |
| 18.5-24.9 (healthy weight) | 1,325 (61.9%) |
| 25.0-29.9 (overweight) | 290 (13.5%) |
| ≥ 30.0 (obese) | 85 (4.0%) |
| Missing | 95 (4.4%) |
| **Underlying conditions** |  |
| Modified Charlson Comorbidity Index - n (%) |  |
| 0 | 1,368 (63.9%) |
| 1 | 328 (15.3%) |
| 2 | 368 (17.2%) |
| 3 | 57 (2.7%) |
| 4-6 | 21 (< 1%) |
| Diabetes - n (%) | 1,433 (66.9%) |
| Lung disease - n (%) | 186 (8.7%) |
| Liver disease - n (%) | 58 (2.7%) |
| Heart disease - n (%) | 94 (4.4%) |
| Renal disease - n (%) | 452 (21.1%) |
| Stroke - n (%) | 53 (2.5%) |
| Cancer - n (%) | 57 (2.7%) |

To screen candidate host and clinical factors for use as covariates, we first performed univariable logistic regression with 28-day mortality as a binary outcome (**Figure 1c**). Most recorded host comorbidities showed significant positive associations with mortality, although diabetes and thalassaemia showed significant inverse associations. Such inverse associations are consistent with established clinical knowledge^17,18,36^, but may also have reflected selection or collider bias arising from recruitment conditional on hospital presentation and culture- confirmed infection^28^, rather than protective effects of these conditions. To reduce reliance on individual comorbidity estimates, we therefore summarised comorbidity burden using a modified Charlson Comorbidity Index (CCI)^37^, with modifications according to risk factors recorded in the patient cohorts (**Supplementary Table 2**). We also observed that age and whether the patient was referred to the study hospital from another facility also showed significant positive associations with mortality and that some study hospitals were significantly less associated with death compared to the reference tertiary hospital.

We next fitted a multivariable logistic regression model incorporating age, sex, BMI, CCI, hospital of admission and time to definitive antibiotic treatment to assess their independent associations with 28-day mortality (**Figure 1c**). Sex, BMI, and time to definitive antibiotic treatment were retained in this model despite the absence of significant univariable associations, based on their epidemiological and clinical relevance. Male sex is associated with occupational exposure to *B. pseudomallei* particularly among agricultural workers exposed to contaminated soil and water through skin wounds^38,39^, while BMI has previously been associated with the risk of developing melioidosis^39^. Time to antibiotic treatment was included to account for potential differences in clinical management that could influence mortality. In the multivariable model, age and comorbidity burden remained independently associated with mortality: each 10-year increase in age was associated with higher mortality (adjusted odds ratio [_adj_OR] = 1.14, 95% confidence interval [CI] = 1.05-1.24), as was each unit increase in CCI (_adj_OR = 1.38, 95% CI = 1.24-1.54). Neither sex nor BMI were independently associated with mortality after adjustment. Time to definitive antibiotic treatment showed a borderline association, with earlier treatment associated with lower odds of death (_adj_OR = 0.961, 95% CI = 0.911-1.01). Hospital of admission was also associated with mortality, with the tertiary hospital in Ubon Ratchathani showing the highest observed odds of death (_adj_OR = 1.84, 95% CI 1.45- 2.32).

To determine the covariates appropriate for estimating bacterial genetic associations with mortality, we constructed a directed acyclic graph (DAG) representing hypothesised relationships between bacterial genetic variation, host characteristics, clinical management and host mortality (**Figure 1d**). Although sex and BMI were not independently associated with 28-day mortality in the multivariable clinical model, we retained both alongside age and CCI as core adjustment variables. This approach accounted for baseline host characteristics that could differ across patients carrying different bacterial variants, without restricting covariate selection solely to variables showing independent associations with mortality. In contrast, hospital of admission and time to definitive antibiotic treatment could lie downstream of bacterial genetic effects through disease severity, referral patterns and clinical management. Adjusting for these variables in the primary analysis could therefore attenuate pathogen-associated effects operating through these pathways. We consequently include age, sex, BMI and CCI as core covariates in the primary bacterial genetic analyses, while evaluating additional adjustment for hospital of admission and time to definitive antibiotic treatment in sensitivity analyses due to their positions as mediators.

### Bacterial population structure and predominantly single-strain infections support bacterial association mapping

Of the 2,142 patients recruited, 2,015 (94.1%) had at least one sequenced *B. pseudomallei* genome, including 49 patients from whom multiple bacterial isolates were sequenced (**Figure 1b, Supplementary Tables 3-4**). This yielded 2,101 *B. pseudomallei* genomes for population genomics analysis, providing, to our knowledge, one of the largest collections of *B. pseudomallei* genomes linked prospectively to individual melioidosis outcomes. We characterised the population structure of the infecting isolates using two complementary approaches: a core-genome phylogeny built from reads mapped to the 1996 reference strain K96243^40^, and a density-based clustering of genome assemblies using PopPUNK^41^ (see **Methods** and **Supplementary Table 4**). The two approaches identified concordant, well- separated clusters, resolving seven major lineages (n > 30 per lineage): lineage 1 (n = 447; including K96243), lineage 2 (n = 363), lineage 3 (n = 232), lineage 4 (n = 155), lineage 5 (n = 128), lineage 6 (n = 43), and lineage 7 (n = 37). Together, these lineages comprise 1,405 of 2,101 isolates (66.9%), with the remaining 696 isolates (33.1%) distributed across minor lineages and singletons (**Figure 2a**). These isolates were retained in downstream analyses rather than excluded to avoid selection bias. The distribution of major lineages was consistent with previous population structure analyses of a subset of this cohort^20^, indicating that the collection captures the dominant *B. pseudomallei* lineages infecting patients in northeast Thailand and is not simply an artefact of the current sampling frame.

**Figure 2:**
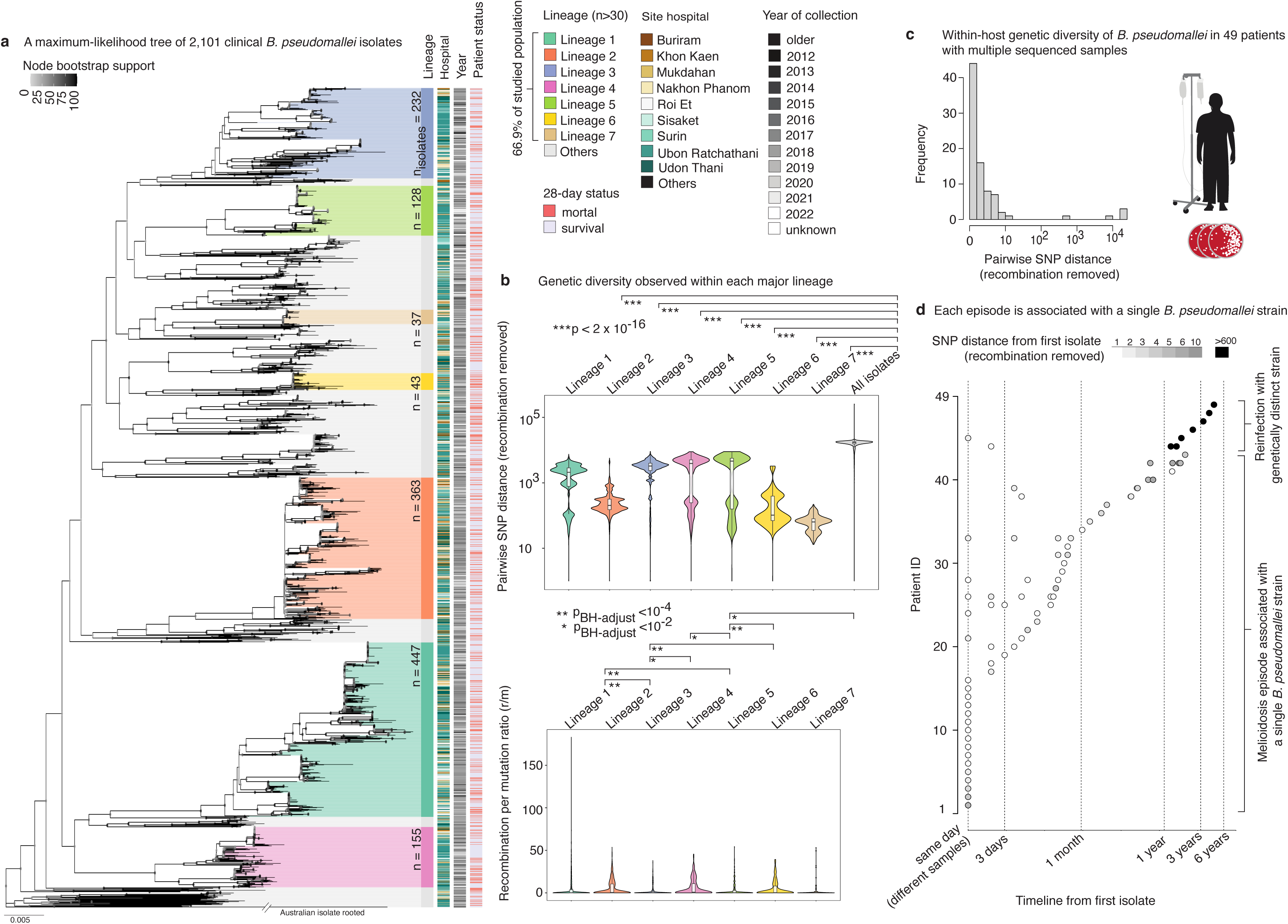
Population structure of clinical *B. pseudomallei* in northeast Thailand and the genetic diversity of infecting strains within a patient. (a) Maximum-likelihood phylogeny of 2,101 clinical *B. pseudomallei* isolates, rooted on an Australian isolate, and bootstrapped 320 times (node bootstrap support). Seven lineages each containing more than 30 isolates are highlighted, and all remaining isolates are grouped as others. The four annotation columns to the right of the tree give, for each isolate, its lineage, the recruiting hospital, the year of collection, and the 28-year outcome of the patient from whom it was cultured (red, died; lilac, survived). **(b)** Genetic diversity within each major lineage. **Top**: distribution of pairwise SNP distance after removal of recombined regions, shown for each seven lineages and for all isolates combined. Every lineage is less diverse than the collection as a whole (p < 2 × 10^-16^, Mann-Whitney test). **Bottom**: distribution of the recombination-to-mutation ratio (r/m) within each lineage. Horizontal bars mark pairwise comparison between lineages that remained significant after Benjamini-Hochberg correction (**p < 10^-4^, *p <10^-2^); comparisons without a bar were not significant. Violins show the full distribution with the median and interquartile range. **(c)** Within-host genetic diversity among the 49 patients from whom more than one isolate was sequenced. The histogram summarises pairwise SNP distance, after removal of recombined regions, between isolates recovered from the same patient. **(d)** Genetic distance between serial isolates and the first isolate from the same patient, plotted against the time elapsed since the first isolate. Each row is one patient (n = 49) and each point one subsequent isolate, shaded by its SNP distance from the first isolate (greyscale for 1-10 SNPs, black for > 600). Isolates recovered within a single episode of melioidosis were near-identical to the first isolate, whereas the distant pairs (black) correspond to reinfection with a genetically distinct strain, in all cases recovered after a prolonged interval.

To provide additional genetic evidence for the lineages, we partitioned genetic variation attributable to recombination as opposed to point mutations using recombination-masked alignments generated with Gubbins^42^, then quantified genetic diversity within and between lineages. Pairwise SNP distances were substantially smaller within lineages than across the population (within-lineage median 938 SNPs, IQR 177–2,905; population-wide median 17,040 SNPs, IQR 15,559–18,210; two-sided Mann-Whitney U test, p < 2 × 10^-16^; **Figure 2b**; **Supplementary Table 5**), confirming that the lineages represented distinct genetic backgrounds. The ratio of recombination to mutation (r/m) also varied among lineages (median 0.269, IQR 0.143-0.429, χ^2^test p = 1.02 × 10^-9^; **Supplementary Table 6**), indicating that in different lineages, recombination contributed to genetic diversity to different degrees.

We then assessed whether melioidosis episodes were typically caused by a single or multiple infecting strains, which is important for linking bacterial genotype to clinical outcome. Among the 49 patients with multiple sequenced isolates, sampling intervals ranged from the same date to 1,752 days. Within-host pairwise SNP distances, calculated after masking recombinant regions, showed a bimodal distribution (**Figure 2c**). Most isolate pairs differed by 0-10 SNPs, whereas a second distinct cluster differed by > 600 SNPs. Given that *B. pseudomallei* has an estimated within-host core-genome mutation rate of ≤ 18.6 SNPs per genome per year^43^ and the isolates’ chronologies, we deduced two infection patterns: (1) isolates differing by 0-10 SNPs were recovered from multiple specimen types over intervals of up to two years, consistent with persistence of a single infection strain, (2) isolates differing by > 600 SNPs recovered more than one year apart were consistent with reinfection by genetically distinct strains (**Figure 2d**). Together, these analyses indicated that individual melioidosis episodes were predominantly associated with a single *B. pseudomallei* isolate. For patients with multiple distinct episodes, only the first episode was retained for the primary association analysis as host clinical data were collected then. This approach thus established a one-to-one bacterial genotype-clinical outcome mapping while minimising potential confounding from within-host persistence or reinfection.

### Host factors and lineage 6 explain mortality variation

Having defined bacterial lineages and observed variation in patient mortality status across lineages (**Figure 2a**), we hypothesised that host factors and bacterial lineage together explain a substantial proportion of mortality variation. We quantified the relative contribution of host and clinical factors identified in our causal framework and of bacterial lineage to mortality using variance decomposition (**Table 2**). CCI explained up to 4.8% of the variation in 28-day mortality (liability-scale R^2^ 0.048, 95% CI 0.021-0.088), followed by age at 4.3% (0.043, 95% CI 0.028-0.087) and hospital of admission at 3.7% (0.0377, 95% CI 0.0212-0.0844). Bacterial lineage (**Supplementary Table 3**) alone explained 2.4% of the variation (0.0239, 95% CI 0.0126-0.0654), whereas the combined model accounted for 15.1% (R^2^ 0.151, 95% CI 0.138-0.245). Consistent with these estimates, adding host characteristics to a model containing bacterial lineage substantially improved model fit (likelihood ratio test [LRT] p = 3.66 × 10^-10^), whereas adding bacterial lineage to the host model produced a more modest improvement (LRT, p = 0.028). Together, these findings showed that host characteristics account for a substantial component of mortality variation and provided a basis for accounting for host heterogeneity in subsequent analyses of bacterial determinants of outcome.

**Table 2.**
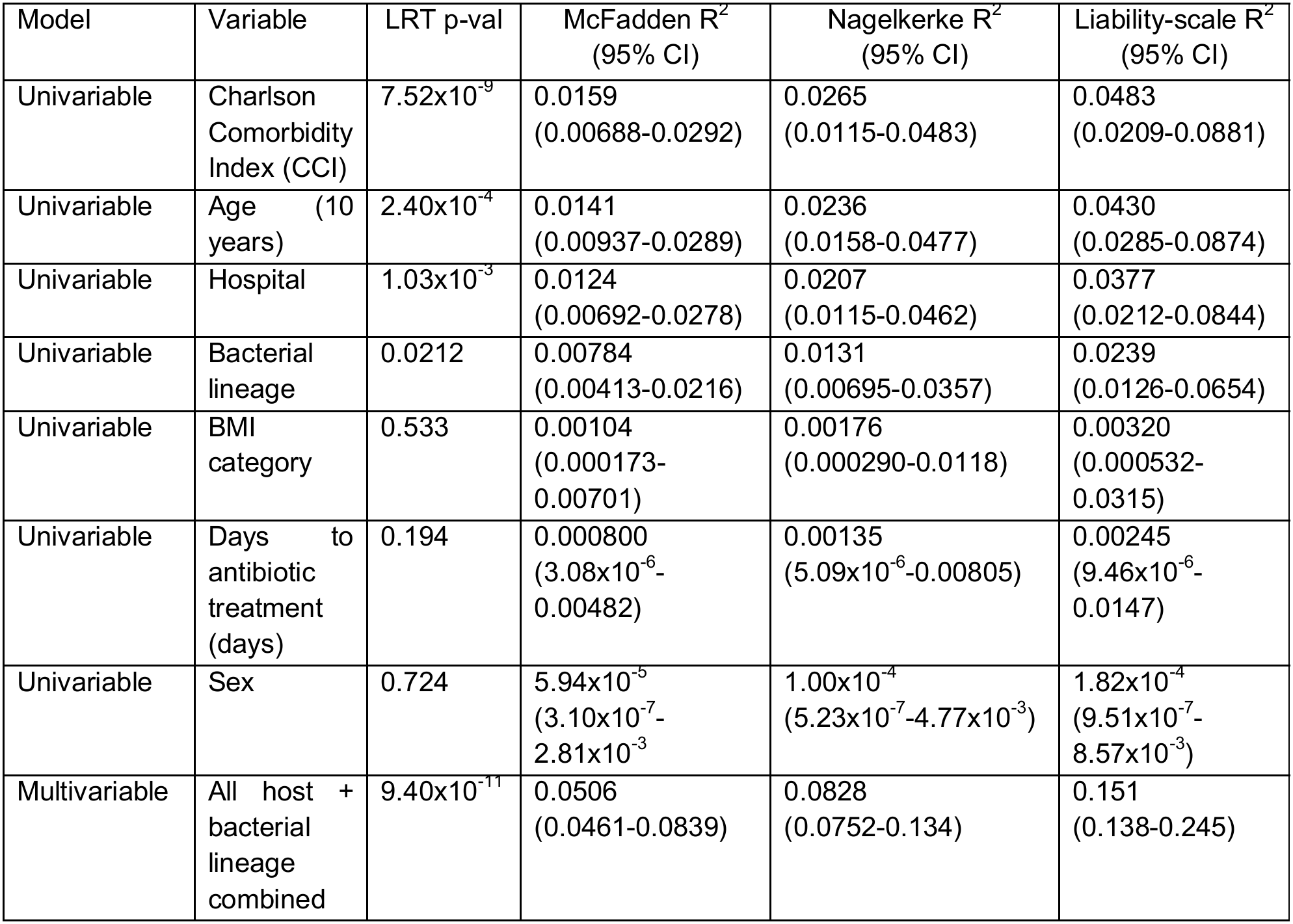
Variance decomposition (n = 1,821; patients with missing data in any field were excluded, mortality prevalence was 26.3%)

| Model | Variable | LRT p-val | McFadden R<sup>2</sup><br>(95% CI) | Nagelkerke R<sup>2</sup><br>(95% CI) | Liability-scale R<sup>2</sup><br>(95% CI) |
|---|---|---|---|---|---|
| Univariable | Charlson Comorbidity Index (CCI) | 7.52x10<sup>-9</sup> | 0.0159<br>(0.00688-0.0292) | 0.0265<br>(0.0115-0.0483) | 0.0483<br>(0.0209-0.0881) |
| Univariable | Age (10 years) | 2.40x10<sup>-4</sup> | 0.0141<br>(0.00937-0.0289) | 0.0236<br>(0.0158-0.0477) | 0.0430<br>(0.0285-0.0874) |
| Univariable | Hospital | 1.03x10<sup>-3</sup> | 0.0124<br>(0.00692-0.0278) | 0.0207<br>(0.0115-0.0462) | 0.0377<br>(0.0212-0.0844) |
| Univariable | Bacterial lineage | 0.0212 | 0.00784<br>(0.00413-0.0216) | 0.0131<br>(0.00695-0.0357) | 0.0239<br>(0.0126-0.0654) |
| Univariable | BMI category | 0.533 | 0.00104<br>(0.000173-0.00701) | 0.00176<br>(0.000290-0.0118) | 0.00320<br>(0.000532-0.0315) |
| Univariable | Days to antibiotic treatment (days) | 0.194 | 0.000800<br>(3.08x10<sup>-6</sup>-0.00482) | 0.00135<br>(5.09x10<sup>-6</sup>-0.00805) | 0.00245<br>(9.46x10<sup>-6</sup>-0.0147) |
| Univariable | Sex | 0.724 | 5.94x10<sup>-5</sup><br>(3.10x10<sup>-7</sup>-2.81x10<sup>-3</sup> | 1.00x10<sup>-4</sup><br>(5.23x10<sup>-7</sup>-4.77x10<sup>-3</sup>) | 1.82x10<sup>-4</sup><br>(9.51x10<sup>-7</sup>-8.57x10<sup>-3</sup>) |
| Multivariable | All host + bacterial lineage combined | 9.40x10<sup>-11</sup> | 0.0506<br>(0.0461-0.0839) | 0.0828<br>(0.0752-0.134) | 0.151<br>(0.138-0.245) |

To investigate the lineages’ association with mortality, we then performed multivariable logistic regression with 28-day mortality as the binary outcome. We used lineage 1 as the reference because it is the most common lineage and includes the K96243 reference strain^40^. We adjusted for host characteristics (age, sex, BMI and CCI) and further performed sensitivity analyses incorporating hospital of admission and time to definitive treatment to assess whether the lineage mortality associations were robust to additional adjustment for clinical factors (**Supplementary Figure 1**). We found that lineage 6 was associated with higher mortality than lineage 1, including after adjusting for host covariates (median host-adjusted OR = 2.646, 95% CI 1.314-5.294, p < 0.007, **Figure 3a**). This association remained consistent across models using different combinations of host and clinical covariates, with neither the likelihood nor the Akaike Information criterion (AIC) and Bayesian information criterion (BIC) clearly favouring a particular covariate combination (**Supplementary Figure 1**). To account for variation in days to death, especially given that deaths occurred mostly early during severe sepsis (**Supplementary Figure 2a**), we additionally performed Cox proportional hazards analyses using days to death as the outcome, till 28 days. Consistent with the logistic regression analysis, the Cox proportional hazards model identified an increased hazards of death among patients infected with lineage 6 compared with lineage 1 (adjusted HR = 2.084, 95% CI 1.255-3.460, BH-adjusted p = 0.0272; **Supplementary Figure 2b**). No other lineage showed a consistent association with mortality across models. These findings identified lineage 6 as a candidate marker of increased mortality risk, with the association robust to adjustment for host characteristics and additional clinical covariates.

**Figure 3:**
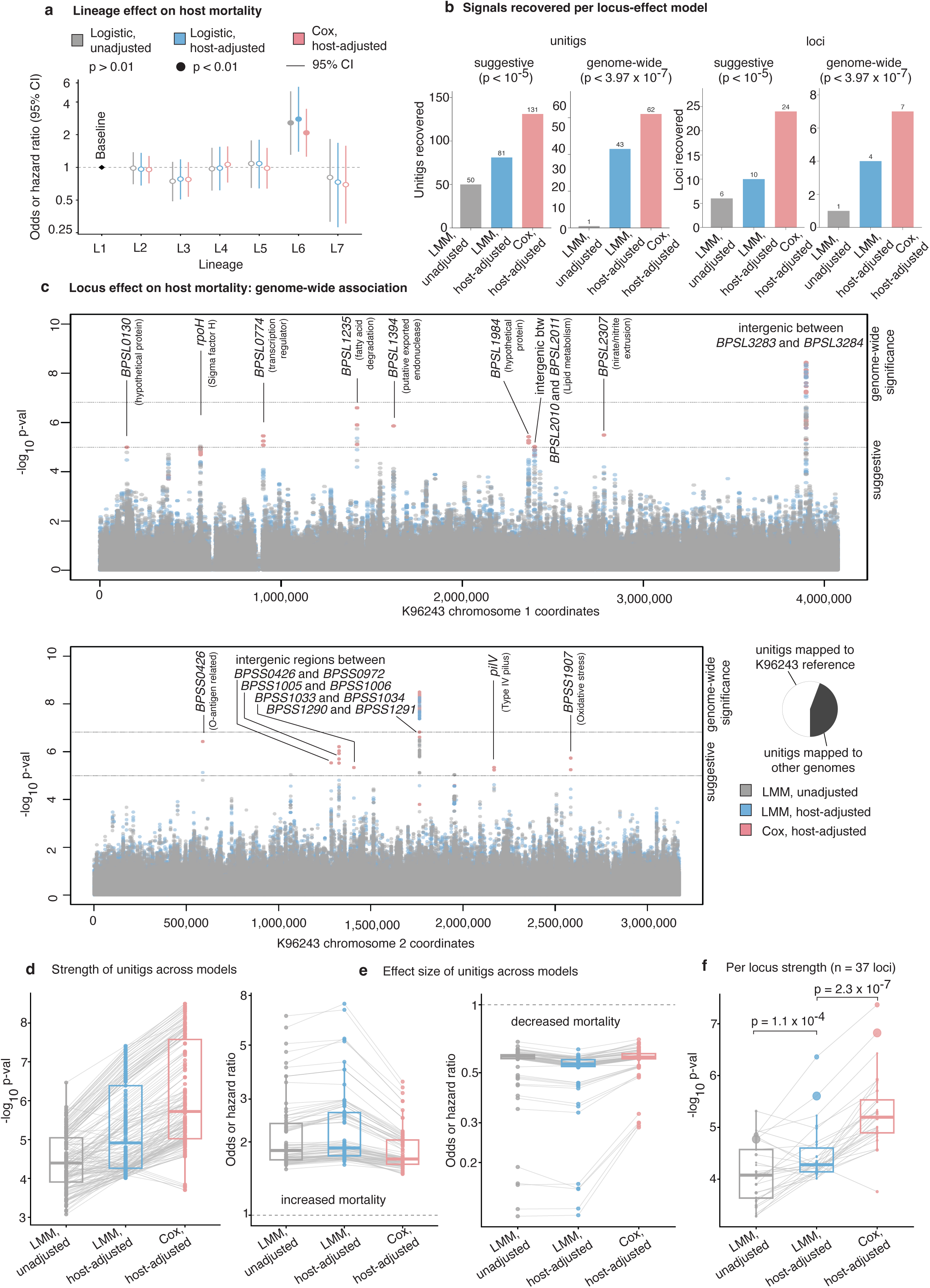
**Host-covariate adjustment increases power to detect bacterial determinants of melioidosis mortality**. **(a)** Association between *B. pseudomallei* lineage and 28-day mortality (n = 2,014 patients, one isolate per patient), with lineage 1 as reference. Points show odds ratios (multivariable logistic regression) or hazard ratios (Cox proportional hazards) and error bars the 95% confidence interval; filled points indicate p < 0.01, open points p > 0.01 (two-sided Wald test). Unadjusted models accounted for lineage alone; host-adjusted models additionally accounted for age, sex, BMI category and Charlson comorbidity index. **(b)** Number of mortality- associated signals recovered by each model as unitigs (left) and independent loci (right), at a suggestive threshold (p < 10^-5^) and the permutation-derived genome-wide threshold (p < 3.97 × 10^-7^; 100 permutations of mortality status on fixed genotypes). **(c)** Genome-wide association with 28-day mortality across chromosome 1 (**top**) and chromosome 2 (**bottom**). Each point is a unitig, coloured by the model giving the strongest association at that position; dashed lines mark the thresholds in (b). **(d, e)** Evidence strength (d) and effect size (e) for top unitigs across models (174 unitigs reaching p < 10^-4^ in the host-adjusted linear mixed model); grey lines connect the same unitig. Boxes show the median and interquartile range, whiskers 1.5× the interquartile range. Effect sizes are shown separately for unitigs associated with increased and decreased mortality. **(f)**, as in (d) with each of 37 independent loci as the unit of analysis (median across constituent unitigs). P values are from two-sided Wilcoxon signed-rank tests on per-locus difference.

### Host adjustment increases the power to detect bacterial loci associated with mortality

After identifying an association between bacterial lineage 6 and mortality, we next asked whether genetic variation at specific bacterial loci explained additional variation in mortality and whether adjustment for host characteristics increased power to detect these finer-scale bacterial associations. We represented bacterial genetic variation using unitigs (see **Methods**) and performed a GWAS of 28-day mortality using a linear mixed model (LMM) with random effect to account for bacterial population structure. The model adjusted for bacterial lineages but unadjusted for host characteristics showed little evidence of genomic inflation (λ_LMM_ = 1.007, **Supplementary Figure 3**). Adjusting for the core host confounders identified in our causality modelling framework (age, sex, BMI and CCI) strengthened bacterial genetic association signals, whereas additional adjustment for the sensitivity variables – hospital of admission and time to antibiotic treatment – attenuated the signals. Acknowledging the hypothetical nature of the relationships identified in the directed acyclic graph (**Figure 1d**), we performed a sensitivity analysis with covariates (**Supplementary Figure 5**), which confirmed age, sex, BMI, and CCI as the optimal set of covariates; therefore, we used these confounders as covariates in subsequent analysis. After adjustment for age, sex, BMI and CCI, the LMM showed reduced genomic inflation (λ_LMM_ = 0.984, **Supplementary Figure 3),** and stronger unitig-level associations with mortality (**Figure 3b-c**). To investigate associations with time to death, we selected unitigs associated with mortality in the host-adjusted LMM at a relaxed threshold (p < 10^-4^) to prioritise relatively high-significance unitigs for further characterisation and then tested them using a mixed-effects Cox model with lineage as a random effect^44^. The Cox analysis provided stronger statistical evidence for associations between the selected unitigs and mortality (**Figure 3b-f**).

Associated unitigs mapped to genes implicated in virulence, stress response, metabolism, and regulation, as well as intergenic regions in the K96243 reference genome. Many unitigs, however, did not map to this reference (**Figure 3c, Supplementary Figure 6**). As multiple associated unitigs frequently mapped to the same genomic regions in the reference (**Figure 3c**) or across isolate assemblies, we grouped them into loci based on overlapping genomic alignments and population frequencies (see **Methods**). This grouping enabled us to assess the distribution of association signals at the locus level and to prioritise candidate regions for further investigation. Host adjustment increased the number of unitigs and loci reaching the predefined suggestive threshold (p < 10^-5^) and permutation-based genome-wide significance (p < 3.97 × 10^-7^) thresholds (see **Methods**). In the unadjusted LMM, 50 unitigs representing six loci reached the suggestive threshold, whereas one unitig representing one locus reached genome-wide significance. After adjustment for age, sex, BMI, and CCI, 81 unitigs representing ten loci reached the suggestive threshold and 43 unitigs representing four loci reached genome-wide significance. The host-adjusted Cox analysis provided further gains: 131 unitigs representing 24 loci reached the suggestive threshold, and 62 unitigs representing seven loci reached genome- wide significance (**Figure 3f**; Wilcoxon test, p < 1.1 x 10^-4^). Despite these differences in statistical significance, effect estimates remained highly correlated between models (**Figure 3e**), suggesting that host adjustment primarily strengthened the statistical evidence for existing associations rather than substantially changing their estimated directions of effect.

### Examples of mortality-associated loci and their lineage-specific distribution

We prioritised 24 loci containing unitigs that met the suggestive significance threshold in the host-adjusted Cox analysis for downstream functional investigation (**Supplementary Table 7**). These loci included genes with established or plausible roles in *B. pseudomallei* biology. For example, *pilV* (ML29) encodes a component of the type IV pilus system which contributes to host colonisation and adhesion, while *BPSL2307* (ML33) encodes a putative nitrite extrusion protein, and is potentially implicated in bacterial adaptation to nitrosative stress under severe hypoxic conditions within the host. Two additional loci (ML23 and ML32) mapped to orthologues of *BPSL0774*, which encodes a sensor kinase involved in two-component signal transduction. Notably, many loci mapped to intergenic regions or hypothetical genes in the accessory genome rather than core annotated genes in K96243 reference. Some of these loci showed stronger associations with mortality than loci mapping to annotated reference genes, highlighting the potential contribution of poorly characterised and accessory genomic regions to variation in disease outcome.

We next investigated whether these mortality-associated loci were distributed broadly across the bacterial population or concentrated in particular lineages or sublineages (**Supplementary Figure 4a**). The distribution of locus-associated unitigs varied across lineages, with some showing lineage-specific patterns consistent with ancestral gene gain or loss. For example, some unitigs at ML5, an intergenic region between *BPSL3283* (encoding a hypothetical protein) and *BPSL3284* (encoding a secreted substrate binding protein) were absent from lineage 5. To assess whether mortality-associated genetic variation was enriched in particular lineages, we compared the distribution of one representative mortality-associated unitig per locus across isolates. Lineage 6, which we previously found associated with mortality, carried significantly more of these unitigs than isolates from other lineages (Kruskal–Wallis test, p < 10^-40^, **Supplementary Figures 4b-c**). Post hoc Dunn’s tests with Benjamini–Hochberg correction showed that the median number of mortality-associated unitigs per isolate was three in lineage 6, compared with 0 to 2 in the other lineages. These population-level patterns indicate that mortality-associated genetic variation was not uniformly distributed across the bacterial population. Although the mixed-model framework accounted for population structure when testing individual genetic associations, this enrichment in lineage 6 highlights the potential contribution of lineage-linked genetic backgrounds to disease outcome and the importance of distinguishing locus-level from broader lineage-level associations.

### Bacterial locus *pilV* is associated with variation in neutrophil populations, including immature compartment

We next asked whether mortality-associated bacterial lineages and loci were associated with distinct circulating host immune cell populations (**Figure 4a**). Among the patients with bacterial genomes sequences, host bulk blood transcriptomes were available from 646 patients across four cohorts. In each cohort, blood was sampled for transcriptomics at different time points after admission: 0 days in DORIM (n = 28) and Ubon-Sepsis (n = 144), approximately 4 days in BurkHostGEN (n = 393), and approximately 5 days in MICRO1501 (n = 81) (**Figure 4b**). Given the expected changes in host immune responses during infection and following antibiotic treatment, cohorts were analysed separately. From the transcriptomes, we imputed the absolute abundance of 23 immune cell types using CIBERSORTx^45^ and tested their association with mortality-associated bacterial lineages and loci using generalised linear models^46^. Analyses were restricted to groups with at least seven patients, reflecting the seven model parameters required to account for the intercept, age, sex, comorbidity burden and BMI categories (**Figure 4c-d**).

**Figure 4:**
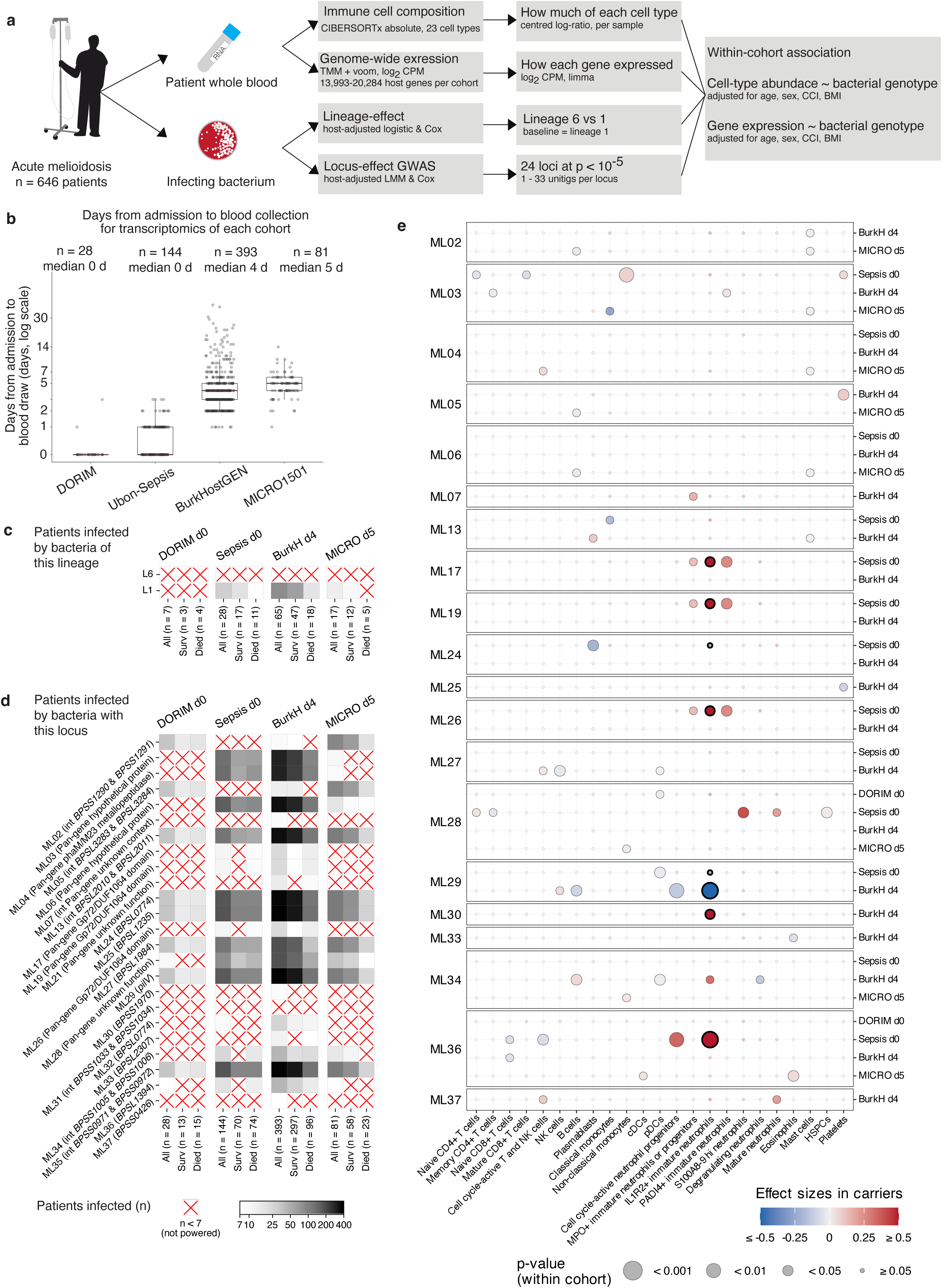
Design and power of the paired host-pathogen analysis and association between bacterial genotype with circulating immune cell composition. **(a)** Analytical workflow. Whole blood and the infecting *B. pseudomallei* isolate were obtained from 646 patients with culture-confirmed melioidosis. Blood transcriptomes were used to estimate the abundance of 23 immune-cell types using CIBERSORTx, expressed as centred log-ratios, and to quantify host gene expression. Bacterial genomes were used to define lineage and locus effects. Of 24 loci associated with 28-day mortality at p < 10^-5^ in 2,014 sequenced isolates, 23 were present in the paired transcriptome cohort and analysed here. **(b)** Interval between hospital admission and blood collection in each cohort. Boxes show the median and interquartile range; whiskers, 1.5 × IQR; points, individual patients. Cohorts are denoted by cohort and sampling day (DORIM d0, Sepsis d0, BurkHostGEN d4 and MICRO1501 d5). **(c)** Number of patients infected with lineage 1 or lineage 6 isolates in each cohort, stratified by 28-day mortality. Cell shading indicates group size; red crosses denote groups with fewer than five patients, excluded from association testing. **(d)** As in (c), for the mortality-associated loci. Locus identifiers are followed by the associated gene or genomic context, or pan-gene for loci outside the K96243 reference. **(e)** Association between bacterial locus carriage and abundance of 23 immune-cell types, tested separately within each adequately powered cohort. Rows are grouped by locus and cohort; columns represent cell types ordered from lymphoid to myeloid lineages. Point colour indicates the estimated difference between carrier and non-carrier isolates in centred log-ratio units (blue, lower; red, higher), and point size indicates p-value. Loci for which all p-values were > 0.05 were not shown. Estimates were adjusted for age, sex, Charlson comorbidity index and body mass index. P-values are unadjusted for multiple testing.

We first examined whether infection with bacterial lineage 6, which was associated with mortality, corresponded to differences in host immune cell composition relative to lineage 1. No cohort contained sufficient lineage 6 infections to meet the prespecified sample-size threshold. In an exploratory analysis of BurkHostGEN, however, we observed a decrease in degranulating neutrophils that remained significant after correction for multiple testing (FDR = 3.65 × 10^-3^). Only four patients in this cohort were infected with lineage 6 isolates, however, precluding reliable inference about a lineage-specific effect on immune cell composition.

We next examined 24 bacterial loci that passed the suggestive significance threshold in the host-adjusted Cox analysis (**Figure 4e**). Of these, 23 were sufficiently prevalent to be tested in at least one transcriptome cohort, yielding 48 locus-cohort pairs and 1,104 tests across the 23 immune cell types. In the largest cohort, BurkHostGEN, the survival-associated ML29 locus, mapping to *pilV,* which encodes a type IV pilus component, was associated with reduced abundance of IL1R2^+^ immature neutrophils (FDR = 7.19 × 10^-3^) and cell cycle-active neutrophil progenitors (FDR = 1.58 × 10^-3^); the locus covers the non-synonymous SNP G201S called by ARIBA^47^ (see **Methods**) situated in an N-terminal constant region. Both associations showed the same direction of effect in another cohort (Ubon-Sepsis) with sufficient sample size, although neither reached statistical significance. To assess whether associations with neutrophils are extended beyond *pilV*, we examined the distribution of nominal associations across the 48 locus-cohort pairs. Seven out of 48 tests reached p < 0.05 for IL1R2^+^ immature neutrophils (binomial p = 4.7 × 10^-4^), suggesting that variation in this neutrophil population might be associated with multiple-mortality associated bacterial loci. This pattern provided hypothesis- generating evidence for a broader relationship between bacterial genetic variation and immature neutrophil abundance, although individual association did not survive correction for multiple testing.

Regulation of neutrophils plays an important role in host defence against *B. pseudomallei*, with both abnormally low and abnormally high neutrophil levels associated with mortality^48^. This is explained by emergency myelopoiesis during sepsis^49^, resulting in high levels of immature neutrophils and proliferating neutrophil progenitors in a shift away from the cell-mediated response^48,50^, or by bone marrow exhaustion or immunoparalysis as the disease becomes more severe^51^. IL1R2^+^ immature neutrophils express IL1R2, a decoy receptor that, when released in soluble form, can sequester the pro-inflammatory cytokine IL-1β, inhibiting IL-1 signalling^52^, which has been shown in mouse models to be deleterious for melioidosis^53,54^. In humans, both IL1R2 and its transcript are predictive of melioidosis mortality^55,56^. Thus, the association between *pilV* variation and reduced abundance of IL1R2^+^ immature neutrophils and cell-cycle active neutrophil progenitors imply maintenance of immature neutrophil populations. Whether these associations reflect altered granulopoiesis, changes in immune cell recruitment, or other host responses remains unresolved.

### Bacterial locus ML34 is associated with coordinated suppression of host blood gene expression

We next investigated whether mortality-associated bacterial lineages and loci were associated with distinct host blood transcriptional responses. We predefined a minimum sample size of 30 patients per bacterial-carriage group for differential expression analysis, based on power considerations for detecting a moderate-to-large expression effect (1.5-fold change) at an FDR < 0.05. Lineage 6 did not reach this threshold in any cohort and was therefore excluded. Only the BurkHostGEN cohort provided sufficient paired transcriptome data for locus-level analysis.

Of the 24 mortality-associated loci passing the suggestive significance threshold in the host- adjusted Cox analysis, eight met the sample-size criterion in BurkHostGEN (**Figure 5a**). For each locus, we tested the association between bacterial infection and host whole-blood gene expression, adjusting for age, sex, BMI and CCI. Among the loci, only ML34 carriage was associated with differential host gene expression: 320 genes (301 down-regulated, 19 up- regulated) were differentially expressed in the patients infected with ML34-carrying isolates (n = 35) compared with patients infected with non-carrier isolates (n = 358). ML34 being the only locus associated with differential host gene expression suggests that most mortality-associated bacterial loci do not produce readily detectable transcriptional changes in circulating blood leukocytes, either because their effects are mediated through other biological pathways or because their transcriptional effects are too modest to resolve at the available sample size. ML34 lies within an intergenic region between *BPSS1005*, which encodes a sodium/hydrogen exchange family protein and the *BPSS1006-BPSS1011* operon, which forms part of a putative biosynthetic gene cluster encoding polyketide synthase machinery (**Supplementary Figure 8a**). The region also contains putative transcription factor binding motifs with a palindromic inverted repeat structure (**Supplementary Figure 8b**), raising the possibility that sequence variation at ML34 affects bacterial transcription regulation, although the mechanism linking ML34 carriage to host transcription and mortality remains unresolved.

**Figure 5:**
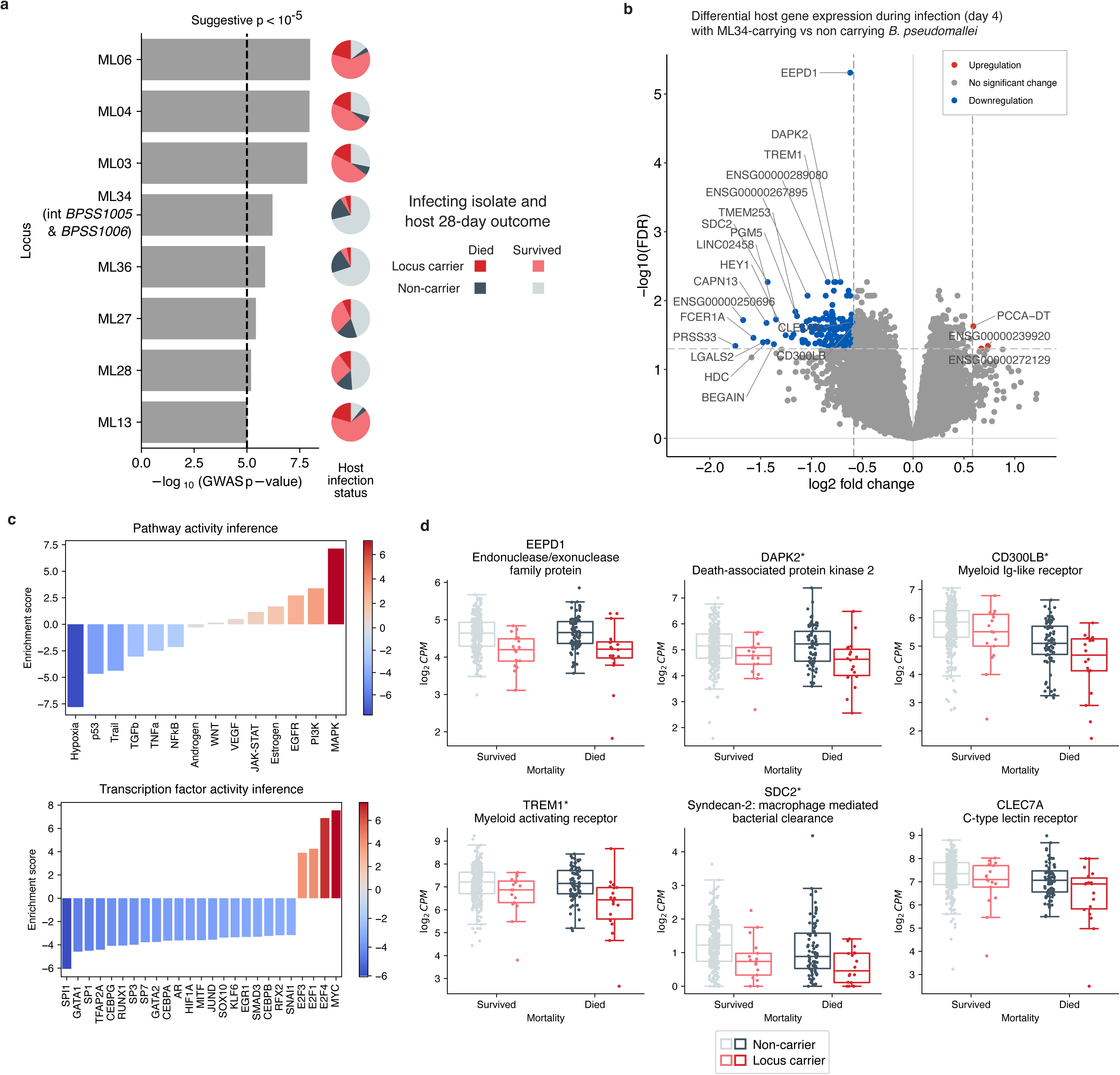
**Carriage of bacterial locus ML34 by infecting B. pseudomallei isolates is associated with down-regulation of host myeloid- and granulocyte-lineage transcripts in BurkHostGEN**. **(a)** Association statistics for eight *B. pseudomallei* loci with suggestive associations with mortality (p < 10^-5^) and carried by at least 30 patients in the BurkHostGEN transcriptome cohort (n = 393). Bars show -log_10_(p) for the lead unitig of each locus from a Cox proportional-hazards model of 28-day survival. Pie charts show the number of patients in the cohort by mortality and carriage of each locus. **(b)** Volcano plot showing differential host gene expression, comparing BurkHostGEN patients infected with ML34-carrying isolates and non- ML34-carrying isolates, controlling for host covariates (age, sex, BMI, CCI) and mortality. Coloured points indicate genes significantly differentially expressed (fold change < 1.5, FDR < 0.05), red indicating up-regulated genes and blue indicating down-regulated genes. **(c)** Pathway and transcription factor activity inference based on differential host gene expression analysis in (b). **(d)** log_2_ counts per million (batch-corrected) for six representative genes across the four carriage-by-outcome groups; boxes show median and interquartile range, whiskers 1.5 × IQR, and points individual patients.

Given that many differentially expressed genes were associated with ML34 carriage and that ML34 carriage was associated with mortality, it is possible that this transcriptional signature merely reflected a maladaptive host response to mortality rather than an adaptive response to a more severe bacterial strain. To account for this, we additionally adjusted for host mortality in differential expression analysis (**Figure 5b**) and found that 164 genes (161 downregulated, 3 upregulated) remained differentially expressed; permutation testing confirmed that this number of differentially expressed genes was unlikely to have arisen by random chance (FDR < 0.05). Collectively, this expression profile suggests increased immune cell growth and proliferation as evidenced by enrichment of MAPK, PI3K, and EGFR pathway activity; downregulation of the p53 pathway; and enrichment of MYC and E2F-family transcription factors (**Figure 5c**). More specifically, we also observed down-regulation of *EEPD1* (endonuclease/exonuclease), *DAPK2* (death-associated protein kinase 2), *CD300LB* (myeloid Ig-like receptor), *TREM1* (triggering receptor expressed on myeloid cells 1), *SDC2* (syndecan-2), and *CLEC7A* (C-type lectin receptor), among other sepsis hallmark genes. Reduced expression of these genes was consistent with altered myeloid activation, and consistent with increased cell growth, rather than a uniformly amplified inflammatory response. These genes also showed lower gene counts with ML34 carriage regardless of mortality status, with mortality only weakly influencing gene expression level (**Figure 5d**), thus suggesting that the transcriptional association with ML34 carriage may not simply reflect the higher mortality observed among carriers.

## Discussion

We found that naturally occurring genetic variation in *B. pseudomallei* was associated with mortality in melioidosis, and that variation at loci was associated with differences in host immune response. Twenty-four loci were associated with mortality at the suggestive significance threshold, of which seven reached genome-wide significance. Two of these could be linked to host transcriptional phenotypes in the same patients: ML29 (*pilV*) was associated with differences in immature neutrophil abundance, while ML34, comprising putative transcription factor binding motifs in the intergenic region upstream of *BPSS1006-BPSS1011*, was associated with differential gene expression of genes involved in myeloid signalling pathways and with pathways implicated in cell proliferation. To our knowledge, this is the first study to associate pathogen genetic variation with both clinical outcome and the host molecular response in the same patient. By integrating bacterial genomic and host transcriptomic data, our findings extend the interpretation of outcome-associated bacterial variants beyond statistical associations towards testable mechanistic hypotheses.

The two loci point to biologically plausible routes to which bacterial variation might influence disease severity. ML29 lies in *pilV*, which encodes a component of type IV pilus apparatus implicated in adherence to host cells. Variation affecting pilus-mediated attachment could possibly alter the magnitude or kinetics of the granulocyte response to infection, consistent with the differences in immature neutrophil abundance observed in patients infected with ML29 genotypes. ML34 lies in an intergenic region upstream of a polyketide synthesis locus. Products of secondary metabolite biosynthetic clusters can have immunomodulatory activity. The association between variation at this locus and downregulation of host myeloid signalling genes is therefore compatible with altered production or regulation of an immunomodulatory metabolite. However, our data cannot distinguish a regulatory effect from the effects of linked variation elsewhere in the cluster. Both interpretations remain hypotheses generated by this study rather than demonstrated mechanisms, but they provide specific directions for experimental investigation.

These findings also have implications for the design and analysis of pathogen genomic studies of clinical outcome. Previous bacterial GWAS have sought to increase statistical power largely through larger sample size^9,12^ and improved control for population structure^57,58^, while adjustment for host characteristics has also been applied in studies of other infections^14^. Our results quantify the potential contribution of host adjustment in a setting with a high burden of comorbidity: the number of loci reaching genome-wide significance increased from one to four after adjustment for host covariates, and to seven when time to death was modelled as a continuous phenotype. Comorbidity burden contributed most to this improvement, consistent with its substantial contribution to variation in mortality in this population. The implications for other infections are likely to be context-dependent, reflecting the extent to which host characteristics explain outcome variation and the specific comorbidity profile of the population studied. Nevertheless, these findings suggest that incorporating host characteristics into pathogen GWAS may improve the identification of bacterial determinants of clinical outcome, particularly in settings where sample sizes are constrained. Integrating host molecular response may further help prioritise candidate loci for mechanistic investigation.

Several limitations should be considered. This observational genomic association study cannot establish causality between bacterial genetic variation and host immune response. Lineage- and locus-level associations should therefore be considered hypothesis-generating pending experimental validation, especially given that estimation of cell proportions from gene expression data gives relative measures with less fidelity for rarer cell types. Transcriptomic data were available for a subset of patients, limiting both lineage- and locus-based analyses. Nevertheless, the reduced abundance of immature neutrophils associated with patients infected with ML29 (*pilV*) across cohorts, together with the downregulation of myeloid-related genes in patients infected with ML34 regardless of mortality, supports a genuine bacterial genotype- associated transcriptional phenotype rather than an artefact. The relatively small transcriptomic sample size, however, warrants caution in interpreting the magnitude and specificity of these effects. Isogenic mutagenesis and complementation of the leading candidate variants, together with experimental studies of the implicated immune pathways, will be required to establish the molecular mechanisms underlying these associations. Finally, our cohorts were predominantly recruited in northeast Thailand, and the generalisability of these findings to *B. pseudomallei* populations in other endemic regions remains to be determined. Nonetheless, to our knowledge, this represents the largest integrated host-pathogen study of a neglected tropical disease to date and provides evidence that combining bacterial genetic variation with host molecular phenotypes can help identify candidate pathogen determinants of clinical outcome in high- burden, resource-limited endemic settings.

## Methods

### Study design and patient recruitment

This study combined clinical, bacterial genomic and host transcriptomic data from prospective melioidosis cohorts in northeast Thailand. This study combined clinical data from 2,142 patients (**Supplementary Table 1**): 78 from a melioidosis relapse study (2012-2014), 174 from the Ubon-sepsis study^29^ (2013-2016), 1,352 from the DORIM study^28^ (2015-2018), 100 from the MICRO-1501 study^30^ (2016-2017), and 438 new patients recruited in the BurkHostGEN study^31^ (2019-2023). All cohorts were recruited at Sunpasitthiprasong Hospital, a tertiary hospital in Ubon Ratchathani, except for DORIM which included hospitals in other provinces in northeast Thailand. The newly recruited cohort was approved by the Sunpasitthiprasong Hospital Ethical Review Board (015/62C) and the Oxford Tropical Research Ethics Committee (OxTREC 25-19). Previously published cohorts were approved by the respective institutional review boards described in the original publications. Written informed consent was obtained from all participants or, where appropriate, their legal representatives.

Clinical data were obtained prospectively using standardised case report forms at the time of enrolment. Demographic variables included age, sex, height and weight. Age at admission was recorded in years. Sex was recorded as male/female per clinical record. Body mass index (BMI) was calculated as weight (kg) divided by height squared (m^2^), using measurements nearest to admission. BMI was categorised as underweight, normal weight, overweight and obese (underweight if less than 18.5 kg m^-2^, overweight if exceeding 25.0 kg m^2^ but not 30.0 kg m^-2^, and obese if exceeding 30.0 kg m^-2^) to allow for potential non-linear associations with mortality. Individual comorbidities, including diabetes mellitus, chronic kidney disease, chronic liver disease, malignancy and other underlying conditions, were recorded where available. Pre- existing comorbidities were summarised using a modified Charlson Comorbidity Index (CCI)^37^, excluding weights contributed by clinical variables not recorded by our case report forms, giving a maximum score of 9 rather than 24 (**Supplementary Table 2**). Hospitals of admission were categorised as the Ubon Ratchathani hospital, at which a plurality of patients was recruited, versus other hospitals. Time to effective antimicrobial treatment was defined as the days between admission and administration of the first antimicrobial agent with *in vitro* activity against *B. pseudomallei*, namely ceftazidime, meropenem or imipenem. The primary clinical endpoint was 28-day all-cause mortality after admission following culture-confirmed melioidosis; patients were thus classified as survivors or non-survivors. Patients that were discharged alive fewer than 28 days after admission were followed up at day 28 to determine mortality status. Clinical variables were assessed for completeness before analysis. The number of observations contributing to each analysis is reported in the corresponding results, figures, and supplementary data.

### Host susceptibility modelling

Candidate covariates were screened using univariable logistic regression (retained at p < 0.05). We modelled continuous variables on their original scale, except for age for which we scaled in units of 10 years, height in units of 20 cm, and weight in units of 20 kg to aid interpretation of odds ratios. For categorical variables, we used the Ubon Ratchathani hospital as the reference for the hospital of admission, female as the reference for sex, and a healthy BMI range as the reference for BMI categories. Multivariable logistic regression retained confounders (age, sex, BMI binned in categories, and CCI) and mediators (hospital and time to melioidosis treatment); this multivariable logistic regression was performed only on patients that had values recorded for all variables (n = 1,930). We constructed a directed acyclic graph (DAG) to formalise the causality linking host characteristics, bacterial genotype, and clinical outcome, identifying confounders and mediators (**Figure 1d**).

### Bacterial isolation, culture and whole-genome sequencing

*B. pseudomallei* was isolated from clinical specimens collected from patients with culture- confirmed melioidosis, as part of standard diagnostic procedures at participating hospitals. For most patients, the clinical specimen was a blood sample and a single colony from a positive culture was subcultured for genomic DNA extraction. Forty-nine patients (further discussed in ‘**Within host diversity analysis**’) contributed more than one isolate, some from multiple clinical specimen types. In all cases, genomic DNA was extracted using standard protocols described in the original cohort publications^28–31^ and processed to prepare sequencing libraries of 150-bp paired-end reads, which were sequenced on the Illumina HiSeq2000 or Illumina NovaSeq platform at the Wellcome Sanger Institute, United Kingdom. Read quality was assessed using FastQC v0.12.1. To identify contaminating or misassigned sequences, taxonomic composition of the reads was assessed using Sylph v0.8.1^59^. Reads passing all quality-control criteria were deposited in the European Nucleotide Archive (**Supplementary Table 4**).

### Read mapping and genome assembly

We mapped quality-filtered reads to the K96243 *B. pseudomallei* reference genome^40^ (accession numbers BX571965 for chromosome 1 and BX571966 for chromosome 2) using Bowtie2 v2.5.1^60^. Then, we called single nucleotide polymorphisms (SNPs) and small insertions/deletions, relative to the reference, using BCFTools mpileup v1.17^61^, masking sites showing mapping quality < 50, read depth < 8, or heterozygous/ambiguous calls as missing data. We excluded four genomes with > 50% missing data (ENA run accessions ERR11449046, ERR11449060, ERR12165279, and ERR11449026) and one genome (ERR11449021) with genome coverage < 80%; the mean read depth across the remaining samples was 70.16 (**Supplementary Table 3**). Core-genome SNPs were extracted and concatenated across all isolates to generate a pseudoalignment using SNP-sites v2.5.1^62^, producing a final alignment comprising 488,127 variable sites.

Quality-trimmed reads were assembled de novo using Shovill v1.0.4 or v1.1.0 as implemented in the AllTheBacteria project^63^, for genomes deposited there; the 374 samples that were not present in AllTheBacteria were assembled separately using Shovill v1.1.0 for consistency. Contigs shorter than 200 bp were excluded. Assembly quality was assessed using QUAST v5.3.0^64^. Four assemblies (ENA run accessions ERR3487812, ERR3488145, ERR3488203, and ERR3488064) had > 1,000 contigs, so they were excluded. Furthermore, PopPUNK v2.6.3^41^ was used to perform additional population-based quality checks (poppunk –qc-db) using the K96243 reference genome as the type isolate. This further identified two genomes (ENA run accession ERR6718158 and ERR11449238) with too many zero distances and they were also excluded. Retained assemblies had a median 182 contigs [IQR 167-197], median N50 of 166,807 [IQR 143,867-187,978], and GC content of 68.15% [IQR 68.19-68.23] (**Supplementary Table 4**), consistent with *B. pseudomallei*.

After we filtered sequences as described above, we further removed duplicated sequences – the same sample sequenced more than once. We thus obtained 2,101 sequences (**Supplementary Table 3**) from 2,015 patients. These sequences included 62 from the melioidosis relapse study, 202 from Ubon-sepsis, 1,274 from DORIM, 116 from MICRO-1501, and 447 sequences from BurkHostGEN.

### Bacterial pan-genome analysis

Retained assemblies were annotated using Bakta v1.11.4^65^ with default settings using the v6.0 database^66^, producing annotated assemblies in GFF3 format. These were then used to construct a pan-genome using Panaroo v1.5.2^67^ using clean mode ‘sensitive’ and a protein family sequence identity threshold of 0.92, as used in ^20^. Genes present in > 99% of isolates were classified as core genes and those present in < 1% are broadly grouped as accessory genes. Our study population had 4,359 core genes, consistent with a previous report based on a subset of our population and 14,424 accessory genes – more than reported previously^20^, highlighting additional genetic diversity captured by combining the cohort collections. The resulting gene presence/absence matrix was used to characterise variation in accessory gene content across the collection.

### Bacterial population structure

Population structure across the 2,101 *B. pseudomallei* isolates was assessed using two complementary methods: a phylogenetic approach based on a core-genome pseudoalignment generated by mapping reads against the *B. pseudomallei* K96243 reference^40^, and a cluster- based analysis based on genome assemblies.

For the phylogenetic approach, we masked known genomic islands^23^, likely introduced by recombination, from the pseudoalignment, because frequent homologous recombination in *B. pseudomallei* can inflate apparent relatedness. Then, using the resulting masked pseudoalignment, we constructed maximum-likelihood trees using IQ-TREE v2.3.6^68^ under a TVM + F + ASC + R6 substitution model, with ascertainment bias correction applied given the exclusion of invariant sites. Branch support was assessed using 320 bootstrap replicates. Trees were rooted using an Australian isolate MSHR5619^19^ (read accession ERR298346, annotation accession GCA_963571805.1) as an outgroup, and visualised alongside isolate metadata (geographical origin and disease outcome) using iTOL v6^69^. Based on this tree, we defined major lineages.

To assess genetic recombination in our population, we used Gubbins v3.3.5^42^ to identify and mask recombination regions for each lineage; this ran on an initial maximum-likelihood tree (RapidNJ, then IQ-TREE for subsequent iterations) built from the pseudoalignment without genomic islands removed. This approach resolves recombination within a lineage with greater sensitivity than genomic-island masking alone, though it is not tractable at the whole-population level given the genetic diversity involved. Regions showing elevated SNP density consistent with the default Gubbins threshold were excluded, and a recombination-filtered SNP alignment was generated for each of the seven main lineages, from which a refined maximum-likelihood phylogeny was reconstructed. The proportion of each lineage’s genome affected by recombination, and the ratio of substitutions introduced by recombination relative to point mutation (r/m) are reported in **Supplementary Table 6**.

For the cluster-based analysis, we used PopPUNK^41^ to characterise population structure across our isolates. We first constructed a PopPUNK database from *de novo* genome assemblies to estimate pairwise core and accessory genome distance for all isolate pairs, using k-mer lengths ranging from 15-31 bp (step size 2) and a maximum core distance of 0.53 to define the search space for distance estimation (*poppunk --create-db --output <DATABASE> --r-files <R_FILES> --min- k 15 --max-k 31 --k-step 2 --max-a-dist 0.53*). We then checked the quality of the database (*poppunk --qc-db*). Finally, we assigned clusters by fitting a hierarchical density-based spatial clustering (HDBSCAN) model to the core-accessory distance distribution (*poppunk --fit-model dbscan*). This initial model was subsequently refined (*poppunk --fit-model refine*) to sharpen cluster boundaries and improve separation between lineages. The final model achieved a network score of 0.9856, a network density of 0.0047, and a transitivity of 0.9902, indicating well-resolved, internally consistent clusters with clear separation between strains. Lineage assignment from PopPUNK were cross-checked against the maximum-likelihood phylogeny for concordance, and against existing *B. pseudomallei* multi-locus sequence typing (MLST)^70^ and clade designations from prior studies^20^.

### Within host diversity analysis

To assess within-host bacterial diversity, we identified patients that provided more than one *B. pseudomallei* sequence, either from different clinical specimens collected during the same disease episode or from isolates collected from different disease episodes. There were 49 patients contributing between two to seven isolates each, spanning collection intervals of 2- 1,752 days. Disease episodes were originally defined following established melioidosis clinical conventions^71^: a positive culture occurring within 12 months of completion of eradication antibiotic therapy for a prior culture-confirmed episode was classified as presumptive relapse and assigned to the same episode, whereas a positive culture occurred more than 12 months after completion therapy was classified as presumptive reinfection and originally assigned to a new, independent episode. This clinical classification was made independently of isolate genotype, with genetic relatedness then used to test, rather than define, whether presumptive relapses and reinfections were consistent with within-host evolution of a single strain or acquisition of a genetically distinct strain. Based on this definition of disease episodes, if a patient had more than one disease episode, only the first was retained as clinical data was collected in the first disease episode for each patient.

For each pair of isolates from the same patient, we calculated pairwise SNP distances from the core-genome pseudoalignment, excluding SNPs falling within recombination regions identified by Gubbins – this avoids inflating apparent diversity with horizontally acquired variation. If isolates from a pair belonged to different lineages, we instead excluded SNPs falling within known genomic islands. We then compared the obtained pairwise SNP distances against the expected accumulation of mutations under a molecular clock^43^. Isolate pairs whose observed SNP distance fell within the range expected under neutral within-host evolution from a single ancestral strain over the sampling interval were classified as representing within-host diversification of a single infecting lineage, consistent with relapse. Isolate pairs separated by far more SNPs than expected under this model or assigned to different lineages by both the maximum-likelihood phylogeny and PopPUNK (see ‘**Bacterial population structure**’, above), were classified as representing reinfection with genetically distinct strains, consistent with the clinical classification above.

For each patient we classified as having within-host pathogen evolution, we chose the isolate from the earliest blood sample if available – or, the earliest tissue sample otherwise – as the representative bacterial genotype for downstream GWAS analyses. This was consistent with our assumption, supported by our observations (**Fig 2d**), that each infection episode is caused by a single bacterial genotype.

### Lineage-level association analysis

To test for association between bacterial lineage and 28-day mortality, we performed logistic regression with 28-day mortality as a binary outcome, encoding lineage as a categorical variable with lineage 1 as a reference as it is the oldest and most populous^20^. This analysis included univariable regression, and multivariable regressions with models each adjusting for all possible combinations of host age, sex, BMI, CCI, hospital of admission, and time to melioidosis antibiotic treatment. Multivariable regressions were only performed on patients providing bacterial isolates that had recorded values for all the above six host covariates (n = 1,821). The models yielded odds ratios, accompanied by Wald tests to determine significance; we corrected for multiple testing across lineages using Benjamini-Hochberg correction. To evaluate model fit, we calculated log-likelihood, Akaike information criterion (AIC), and Bayesian information criterion (BIC) for each model. As additional analysis, we performed Cox proportional hazards regression adjusting for age, sex, BMI, and CCI identified based on host susceptibility modelling (see “**Host susceptibility modelling**”).

### Host-adjusted bacterial GWAS

Bacterial GWASes were performed using unitigs derived from quality-filtered genome assemblies as the genetic variant set, capturing SNPs, indels, and accessory-genome variation not represented by core-genome SNPs alone. Association testing was performed using Pyseer v1.4.0^72^, with 28-day mortality modelled as a binary outcome (*pyseer --phenotypes <PHENOTYPE matrix> --covariates <COVARIATE matrix> --use-covariates <FLAGS> --kmers <UNITIGS> -- uncompressed --lmm --lineage --lineage-clusters <LINEAGE definitions> --lineage-file <LINEAGE effects output> --distances <MASH distance matrix> --similarity <SIMILARITY matrix> --min-af 0.01 --max-af 0.99 --max-dimensions 6*). Unitigs present in 1% to 99% of isolates, totalling 426,409 unitigs, were each tested for association with host mortality using a linear mixed model (LMM)^57^. Here, Pyseer fitted unitig presence as a fixed effect and introduced a random effect term to account for genome-wide relatedness among isolates, thus allowing most genetic relatedness structure to be absorbed by random effects while retaining power to detect locus-specific associations. Bacterial population structure was accounted for using a similarity matrix computed based on unitig presence in each assembly.

Genomic inflation was assessed using quantile-quantile plots of observed versus expected - log_10_(*p*) as a test statistic (**Supplementary Figure 3**), and the genomic inflation factor (λ) was reported for each model. To determine a genome-wide significance threshold, we performed permutation testing by randomising mortality labels across the isolates 100 times. We also defined a suggestive association threshold of p < 10^-5^ to capture unitigs with evidence of association falling below genome-wide significance, consistent with conventional GWAS reporting practice.

To show how accounting for host factors affect bacterial GWAS, we performed a sensitivity analysis by running separate models with all possible combinations of age, sex, BMI, CCI, hospital of admission, and time to antibiotic administration as covariates in addition to bacterial population structure (**Supplementary Figure 5**). These analyses shared the same set of patients for which all these covariate values were defined (1,821 patients) to enable comparison. However, once we identified the optimal combination of covariates from this sensitivity analysis (i.e. age, sex, BMI, and CCI), we re-ran models adjusting for bacterial population structure only and adjusting for the optimal set of covariates on different sets of patients, defined by which patients have covariate values defined for each respective set of covariates (**Figure 3**). As additional analysis, we chose unitigs exceeding p < 10^-4^ in the optimal host-adjusted model for Cox proportional hazards regression^44^, modelling time-to-death (till 28 days) using the same set of optimal host covariates.

### Grouping mortality-associated unitigs into genomic loci

To avoid reporting multiple overlapping unitigs representing the same genomic region, we grouped mortality-associated unitigs into genomic loci based on their positions across 2,014 GWAS genomes. Each unitig was aligned to the de novo assembly of each genome using BLAT v36.0, retaining the default hit and recording contig, coordinates, strand and alignment statistics. Unitigs mapping to multiple positions within a genome were classified as multi-copy. Within each genome, overlapping or adjacent unitig alignments on the same contig were merged into per-genome loci using transitive merging. Master loci were then defined as connected components linking unitigs that occurred in the same per-genome locus in at least one genome. Multi-copy occurrences were excluded from linking to avoid joining unrelated regions through repeats; unitigs that were multi-copy in all genomes remained unassigned.

One locus showed distinct unitig frequency bands and frequent proximity to contig ends, consistent with assembly fragmentation, so we manually investigated this locus. We resolved this locus by performing hierarchical clustering of unitigs based on pairwise co-location, defined as the proportion of genomes carrying both unitigs in the same per-genome locus. Average- linkage clustering of 1 − co-location, cut at 0.5, identified three sub-loci, so we replaced this original locus with these three. This process thus grouped 174 unitigs into 37 loci, assigning 1 to 33 unitigs per locus. For subsequent analysis, we used the presence of the unitig in each locus with the strongest association with mortality (smallest p value) in the host-adjusted Cox GWAS, as the lead unitig, as a proxy for locus presence to account for mixed directionalities of association.

To determine whether mortality-associated genetic variation was enriched within specified lineages, mortality-associated unitigs identified from the host-adjusted GWAS (p < 10^-4^) were mapped back onto each bacterial assemblies, and the presence/absence of each unitig was recorded for every isolate. To test whether mortality-associated variation was disproportionality concentrated within particular lineages, we compared the mean number of mortality-associated major loci (representative unitig has a positive effect size for association with mortality) present per isolate against the mean number of survival-associated major loci (representative unitig has a negative effect size for association with mortality). This comparison also accounted for directionality of association. Differences in per-isolate unitig burden across seven lineages were assessed using Kruskal-Wallis test, with post-hoc pairwise comparison (e.g. lineage 6 versus all other lineages) performed using Dunn’s test with Benjamini-Hochberg correction.

### Conversion of unitig-mapped genes to ARIBA-called variants

We called sequence variants of genes containing mortality-associated unitigs (LMM optimal host-adjusted, p < 10^-4^) from raw sequencing reads against the genes’ sequences in the K96243 reference using ARIBA v2.14.6^47^. This process converts the unitig-level association signal – which does not specify the precise position of the variant within the length of the unitig – into an allele-level variant call – thus resolving the signal into a defined amino acid substitution, insertion, or deletion. However, ARIBA-based variant calling relies on assembling short reads against a reference sequence for the gene of interest; therefore, its accuracy is reduced for genes containing repetitive sequences. For genes with repetitive architecture, ARIBA calls were manually inspected for local assembly contigs and variants falling within regions of low mapping confidence were flagged and interpreted with caution rather than reported as high-confidence allele calls. These calls are reported as annotations for the mortality-associated unitigs in **Supplementary Table 7**.

### Functional annotation of mortality-associated loci

Mortality-associated unitigs identified from the host-adjusted GWAS at p < 10^-2^ were mapped back to the *B. pseudomallei* K96243 reference genome using Pyseer’s *annotate_hits_pyseer* script to assign each unitig to its corresponding genomic location and, where applicable, the overlapping annotated gene. Unitigs mapping within an annotated coding sequence were assigned to that gene. Unitigs mapping outside annotated coding sequences were classified as intergenic and annotated according to their position relative to the nearest flanking genes (upstream, downstream, or within a predicted regulatory/untranslated region). The proportion of mortality-associated unitigs falling within intergenic regions was compared against the expected genome-wide proportion of intergenic sequence (∼12.5% of the genome) using a chi-squared test, to assess enrichment or depletion relative to coding genome. To capture variation not present in the reference genome (i.e. accessory genomes), unitigs failing to map to K96243 were mapped against the pan-genome using the same approach; failing that, existing *B. pseudomallei* promoter/regulatory element predictions^73^.

To further contextualise the relationship between genes that mortality-associated unitigs map to, we computed whether pairs of genes were in operon distance (7,000 bp) for visualisation in a network diagram (**Supplementary Figure 6**). For the pan-genes, this computation was based on the number of assemblies that had each pair of pan-genes within 7,000 bp on the same contig, relative to the number of assemblies that had the pair of pan-genes anywhere in the assembly.

### Host whole-blood transcriptome sequencing

Whole blood samples were collected as part of the four cohorts described in ^30,31,56,74^ using Tempus^TM^ RNA blood tubes (Applied Biosystems, Thermo Fisher Scientific), which stabilised RNA immediately upon collection, and stored at -80°C until processing. RNA was extracted using the Tempus^TM^ Spin RNA isolation kit (Invitrogen, Thermo Fisher Scientific), with DNA contamination removed using TURBO^TM^ DNAse Treatment and Removal Reagents. RNA quality and concentration were assessed using microfluidic chip gel electrophoresis (Bioanalyzer 2100, Agilent). Extracted RNA underwent globin depletion and poly(A) enrichment to enrich for mRNA, and libraries were sequenced at the Wellcome Sanger Institute on the Illumina Novaseq platform, generating paired-end 100 bp reads, with a mean library size of approximately 37.8 million reads per sample. Sequencing reads were aligned to the human reference genome (GRCh38) using two-pass STAR aligner (v2.7.3a)^75^), following a standard Nextflow bulk RNA- seq mapping pipeline. Read quality was assessed using FastQC v0.12.1. Samples with low proportions of uniquely mapped reads or with sex-chromosome expression inconsistent with recorded patient sex were excluded as a quality-control step. In sum, good-quality whole-blood transcriptomes were available for 674 patients (**Supplementary Table 1**). Of these patients, 646 also had sequenced bacterial isolates – 28 from DORIM^56^, 144 from Ubon-sepsis^74^, 81 from MICRO-1501^30^, and 393 from BurkHostGEN – and these were used in further analysis.

### Host transcriptomic association with mortality-associated bacterial variants

For each cohort with whole-blood transcriptomes, gene expression was quantified using featureCounts v2.0.0^76^. Lowly expressed genes were filtered out, retaining only genes with > 0.5 counts per million (CPM) in 5% of samples. The filtered matrix was normalised using the trimmed mean of M-values (TMM) method and log2-transformed to generate a CPM expression matrix, using edgeR v4.10.5^77^. The matrix was then batch-corrected using ComBat-seq from SVA v3.60.0^78^.

To investigate whether mortality-associated bacterial loci were associated with differential host gene expression, we defined patient groups by presence/absence of a major locus represented by the presence/absence of the unitig with the strongest association with mortality. We used limma v3.68.5^79^ and voom^80^ to perform differential gene expression analysis based on the filtered – but not batch-corrected – gene expression matrix, fitting a linear model (*host gene expression ∼ 0 + locus presence + age + sex + CCI + BMI + sequencing batch* or *host gene expression ∼ 0 + locus presence + age + sex + CCI + BMI + mortality + sequencing batch*) to each gene. The models included host covariates to account for host-derived variation in gene expression independent of bacterial genotype and accounted for technical variation across sequencing batches; controlling for mortality additionally accounted for how bacterial loci were associated with mortality. DORIM and Ubon-sepsis transcriptomes were each sequenced in a single batch, so for these cohorts, sequencing batch was not incorporated in the model. p- values were adjusted using the Benjamini-Hochberg procedure. A host gene was considered differentially expressed between variant-carrying and non-carrying patients if it had an FDR ≤ 0.05 and absolute fold change ≥ 1.5. Based on this differential expression analysis, we then used Decoupler v1.8.0^81^ to infer pathway activity using PROGENy^82^ pathway signatures and to infer transcriptional factor activity using the CollecTRI database^83^.

### Immune-cell association with mortality-associated bacterial variants

We imputed the abundance of circulating immune cell population from bulk whole-blood transcriptomes using CIBERSORTx^45^ separately for each cohort. The filtered and batch- corrected gene expression matrix obtained in “**Host transcriptomic association with mortality-associated bacterial variants**” was batch-corrected using CIBERSORTx because it did not have a feature to account for sequencing batches. To estimate cell type proportions using CIBERSORTx, we used absolute mode and a reference signature matrix^50^ comprising 23 immune-cell populations.

To investigate whether mortality-associated bacterial lineages and loci were associated with differential host gene expression, we defined patient groups by lineage of the infecting isolate or presence/absence of a major locus represented by the presence/absence of the unitig with the strongest association with mortality. We fitted a generalised linear model (*cell type abundance ∼ 0 + lineage or locus presence + age + sex + CCI + BMI*) to each cell type, then adjusted p- values using the Benjamini-Hochberg procedure.

### Inferring transcription factor binding sites

To infer transcription factor binding sites in the intergenic region major locus 34 is in, we used FIMO^84^ from the MEME suite v5.5.9 to search for occurrences of motifs catalogued in the PRODORIC database^85^ release 2021.9. We used a p-value cut-off of 10^-5^ to filter probable matches. Once the putative binding sites were identified, we compared the sites’ respective transcription factors as described by PRODORIC with *B. pseudomallei* K96243 protein sequences using BLASTP and obtained percent homology values.

## Data availability

Data are available upon request.

## Code availability

Code is available upon request.

## Author contributions

AW: Methodology, software, formal analysis, investigation, data curation, writing – original draft, writing – review & editing.

PB: Methodology, software, formal analysis, investigation, data curation.

KLB: Methodology, software, investigation, data curation, writing – review & editing.

CCho: Methodology, software, formal analysis, investigation, data curation. KA: Investigation.

GW: Investigation. TN: Data curation. DL: Resources.

TB: Investigation.

TDC: Methodology, resources. RP: Resources.

TY: Resources.

PR: Resources.

WL: Methodology.

BK: Supervision.

JK: Conceptualisation, methodology. JC: Conceptualisation, methodology. SJP: Conceptualisation, resources. JP: Conceptualisation, methodology. NPJD: Conceptualisation, resources. SJD: Conceptualisation, resources. TEW: Conceptualisation, resources. NC: Conceptualisation, resources.

NRT: Conceptualisation, methodology.

EED: Conceptualisation, methodology, investigation, data curation, supervision, writing – review & editing.

CChe: Conceptualisation, funding acquisition, methodology, software, formal analysis, visualisation, investigation, resources, data curation, writing – original draft, writing – review & editing, supervision, project administration.

All authors reviewed and approved the final manuscript.

## Acknowledgements

We thank Dr Katika Akksilp and Professor Yot Teerawattananon for their early input on factors associated with patient mortality, which helped shape the direction of this study.

AW was supported by the Wellcome-funded MORU-OUCRU Discovery Research Academy (MODRA) seed award (grant ID 228179/Z/23/Z), PB by a Sanger PhD studentship, and CCho by a Prince Mahidol PhD studentship. CChe was supported by a Wellcome International Intermediate Fellowship (216457/Z/19/Z), a Sanger International Fellowship, and a Wellcome Career Development Award (323077/Z/24/Z). This research was funded in part by the Wellcome Trust (grants 315982 and 206194). The funders had no role in the study design, data collection and analysis, decision to publish, or preparation of the manuscript. For the purpose of Open Access, the author has applied a CC BY public copyright licence to any Author Accepted Manuscript version arising from this submission.

## Notes

### Competing Interest Statement

The authors have declared no competing interest.

## References

1. Kwok, A. J., Mentzer, A. & Knight, J. C. Host genetics and infectious disease: new tools, insights and translational opportunities. Nat. Rev. Genet. 22, 137–153 (2021).

2. Casanova, J.-L. & Abel, L. From rare disorders of immunity to common determinants of infection: Following the mechanistic thread. Cell 185, 3086–3103 (2022).

3. Tonkin-Hill, G., Ruis, C., Bentley, S. D., Lythgoe, K. A. & Bryant, J. M. Within-host bacterial evolution and the emergence of pathogenicity. Nat. Microbiol. 10, 1829–1840 (2025).

4. Barber, M. F. & Fitzgerald, J. R. Mechanisms of host adaptation by bacterial pathogens. FEMS Microbiol. Rev. 48, fuae019 (2024).

5. Power, R. A., Parkhill, J. & de Oliveira, T. Microbial genome-wide association studies: lessons from human GWAS. Nat. Rev. Genet. 18, 41–50 (2017).

6. Chewapreecha, C. et al. Comprehensive Identification of Single Nucleotide Polymorphisms Associated with Beta-lactam Resistance within Pneumococcal Mosaic Genes. PLOS Genet. 10, e1004547 (2014).

7. Coll, F. et al. Genome-wide analysis of multi- and extensively drug-resistant Mycobacterium tuberculosis. Nat. Genet. 50, 307–316 (2018).

8. Farhat, M. R. et al. GWAS for quantitative resistance phenotypes in Mycobacterium tuberculosis reveals resistance genes and regulatory regions. Nat. Commun. 10, 2128 (2019).

9. Kremer, P. H. et al. Pneumococcal genetic variability in age-dependent bacterial carriage. eLife 11, e69244 (2022).

10. Sheppard, S. K. et al. Genome-wide association study identifies vitamin B5 biosynthesis as a host specificity factor in Campylobacter. Proc. Natl. Acad. Sci. 110, 11923–11927 (2013).

11. Chaguza, C. et al. Bacterial genome-wide association study of hyper-virulent pneumococcal serotype 1 identifies genetic variation associated with neurotropism. *Commun*. Biol. 3, 1–12 (2020).

12. Lees, J. A. et al. Joint sequencing of human and pathogen genomes reveals the genetics of pneumococcal meningitis. Nat. Commun. 10, 2176 (2019).

13. Kremer, P. H. C. et al. Genetic Variation in Neisseria meningitidis Does Not Influence Disease Severity in Meningococcal Meningitis. Front. Med. 7, (2020).

14. Goig, G. A. et al. Clinical and bacterial determinants of unfavorable tuberculosis treatment outcomes: an observational study in Georgia. Genome Med. 17, 143 (2025).

15. Meumann, E. M., Limmathurotsakul, D., Dunachie, S. J., Wiersinga, W. J. & Currie, B. J. Burkholderia pseudomallei and melioidosis. Nat. Rev. Microbiol. 1–15 (2023) doi:10.1038/s41579-023-00972-5.

16. Limmathurotsakul, D. et al. Predicted global distribution of Burkholderia pseudomallei and burden of melioidosis. Nat. Microbiol. 1, 1–5 (2016).

17. Wiersinga, W. J. et al. Melioidosis. Nat. Rev. Dis. Primer 4, 1–22 (2018).

18. Currie, B. J. et al. Melioidosis epidemiology and risk factors from a prospective whole- population study in northern Australia. Trop. Med. Int. Health 9, 1167–1174 (2004).

19. Chewapreecha, C. et al. Global and regional dissemination and evolution of Burkholderia pseudomallei. Nat. Microbiol. 2, 1–8 (2017).

20. Seng, R. et al. Genetic diversity, determinants, and dissemination of Burkholderia pseudomallei lineages implicated in melioidosis in Northeast Thailand. Nat. Commun. 15, 5699 (2024).

21. Campos, C. G., Byrd, M. S. & Cotter, P. A. Functional Characterization of Burkholderia pseudomallei Trimeric Autotransporters. Infect. Immun. 81, 2788–2799 (2013).

22. Adler, N. R. L. et al. Systematic Mutagenesis of Genes Encoding Predicted Autotransported Proteins of Burkholderia pseudomallei Identifies Factors Mediating Virulence in Mice, Net Intracellular Replication and a Novel Protein Conferring Serum Resistance. PLOS ONE 10, e0121271 (2015).

23. Tuanyok, A. et al. Genomic islands from five strains of Burkholderia pseudomallei. BMC Genomics 9, 566 (2008).

24. Bzdyl, N. M., Moran, Clare L., Bendo, Justine & Sarkar-Tyson, M. Pathogenicity and virulence of Burkholderia pseudomallei. Virulence 13, 2139063 (2022).

25. Cloutier, M., Muru, K., Ravicoularamin, G. & Gauthier, C. Polysaccharides from Burkholderia species as targets for vaccine development, immunomodulation and chemical synthesis. Nat. Prod. Rep. 35, 1251–1293 (2018).

26. Vander Broek, C. W. & Stevens, J. M. Type III Secretion in the Melioidosis Pathogen Burkholderia pseudomallei. Front. Cell. Infect. Microbiol. 7, (2017).

27. French, C. T. et al. Virulence from the rhizosphere: ecology and evolution of *Burkholderia pseudomallei*-complex species. Curr. Opin. Microbiol. 54, 18–32 (2020).

28. Chantratita, N. et al. Characteristics and one year outcomes of melioidosis patients in Northeastern Thailand: a prospective, multicenter cohort study. Lancet Reg. Health - Southeast Asia 9, (2023).

29. Hantrakun, V. et al. Clinical epidemiology and outcomes of community acquired infection and sepsis among hospitalized patients in a resource limited setting in Northeast Thailand: A prospective observational study (Ubon-sepsis). PLOS ONE 13, e0204509 (2018).

30. Rongkard, P. et al. Diabetes mellitus is associated with a shared hyper-inflammatory immune response in melioidosis and tuberculosis patients: an observational case-control study. 2024.05.31.24308267 Preprint at 10.1101/2024.05.31.24308267 (2024).

31. Angchagun, K. et al. BurkHostGEN: a study protocol for evaluating variations in the Burkholderia pseudomallei and host genomes associated with melioidosis infection. Wellcome Open Res. 8, 347 (2023).

32. Limmathurotsakul, D., et al. Activities of Daily Living Associated with Acquisition of Melioidosis in Northeast Thailand: A Matched Case-Control Study. PLoS Negl. Trop. Dis. 7, e2072 (2013).

33. Aekplakorn, W., Puckcharern, H., ไทยกลา, ก. & Satheannoppakao, W. Thai National Health Examination Survey V (2014). https://kb.hsri.or.th/dspace/handle/11228/4626 (2016).

34. Aekplakorn, W., Puckcharern, H. & Satheannoppakao, W. Thai National Health Examination Survey VI (2019-2020). https://kb.hsri.or.th/dspace/handle/11228/5425 (2021).

35. Patanavanich, R. & Aekplakorn, W. Thai National Health Examination Survey VII (2024- 2025). https://kb.hsri.or.th/dspace/handle/11228/6360 (2025).

36. Currie, B. J. et al. The Darwin Prospective Melioidosis Study: a 30-year prospective, observational investigation. Lancet Infect. Dis. 21, 1737–1746 (2021).

37. Quan, H. et al. Updating and Validating the Charlson Comorbidity Index and Score for Risk Adjustment in Hospital Discharge Abstracts Using Data From 6 Countries. Am. J. Epidemiol. 173, 676–682 (2011).

38. Cheng, A. C. & Currie, B. J. Melioidosis: Epidemiology, Pathophysiology, and Management. Clin. Microbiol. Rev. 18, 383–416 (2005).

39. Pakdeerat, S. et al. CRISPR-based environmental detection of Burkholderia pseudomallei identifies sanitation gaps and melioidosis risk in northeast Thailand. Nat. Commun. 10.1038/s41467-026-73286-8 (2026) doi:10.1038/s41467-026-73286-8.

40. Holden, M. T. G. et al. Genomic plasticity of the causative agent of melioidosis, Burkholderia pseudomallei. Proc. Natl. Acad. Sci. 101, 14240–14245 (2004).

41. Lees, J. A. et al. Fast and flexible bacterial genomic epidemiology with PopPUNK. Genome Res. 29, 304–316 (2019).

42. Croucher, N. J. et al. Rapid phylogenetic analysis of large samples of recombinant bacterial whole genome sequences using Gubbins. Nucleic Acids Res. 43, e15 (2015).

43. Seng, R. et al. Phenotypic and genetic alterations of Burkholderia pseudomallei in patients during relapse and persistent infections. Front. Microbiol. 14, (2023).

44. Hughey, J. J. et al. Cox regression increases power to detect genotype-phenotype associations in genomic studies using the electronic health record. BMC Genomics 20, 805 (2019).

45. Newman, A. M. et al. Determining cell type abundance and expression from bulk tissues with digital cytometry. Nat. Biotechnol. 37, 773–782 (2019).

46. Zhang, P. et al. Context-specific regulatory genetic variation in MTOR dampens neutrophil- T cell crosstalk in pneumonia-associated sepsis. Nat. Commun. 10.1038/s41467-026-69919-7 (2026) doi: 10.1038/s41467-026-69919-7.

47. Hunt, M. et al. ARIBA: rapid antimicrobial resistance genotyping directly from sequencing reads. *Microb*. Genomics 3, e000131 (2017).

48. Jenjaroen, K., et al. T-Cell Responses Are Associated with Survival in Acute Melioidosis Patients. PLoS Negl. Trop. Dis. 9, e0004152 (2015).

49. Manz, M. G. & Boettcher, S. Emergency granulopoiesis. Nat. Rev. Immunol. 14, 302–314 (2014).

50. Kwok, A. J. et al. Neutrophils and emergency granulopoiesis drive immune suppression and an extreme response endotype during sepsis. Nat. Immunol. 24, 767–779 (2023).

51. Brown, K. A. et al. Neutrophils in development of multiple organ failure in sepsis. The Lancet 368, 157–169 (2006).

52. Peters, V. A., Joesting, J. J. & Freund, G. G. IL-1 receptor 2 (IL-1R2) and its role in immune regulation. Brain. Behav. Immun. 32, 1–8 (2013).

53. Weehuizen, T. A. F. et al. Therapeutic Administration of a Monoclonal Anti-Il-1β Antibody Protects Against Experimental Melioidosis. Shock Augusta Ga 46, 566–574 (2016).

54. Ceballos-Olvera, I., Sahoo, M., Miller, M. A., Barrio, L. del & Re, F. Inflammasome- dependent Pyroptosis and IL-18 Protect against Burkholderia pseudomallei Lung Infection while IL-1β Is Deleterious. PLOS Pathog. 7, e1002452 (2011).

55. Kaewarpai, T. et al. Longitudinal profiling of plasma cytokines in melioidosis and their association with mortality: a prospective cohort study. Clin. Microbiol. Infect. Off. Publ. Eur. Soc. Clin. Microbiol. Infect. Dis. 26, 783.e1–783.e8 (2020).

56. Yimthin, T. et al. Blood transcriptomics to characterize key biological pathways and identify biomarkers for predicting mortality in melioidosis. Emerg. Microbes Infect. 10, 8–18 (2021).

57. Earle, S. G. et al. Identifying lineage effects when controlling for population structure improves power in bacterial association studies. Nat. Microbiol. 1, 16041 (2016).

58. Lees, J. A. et al. Genome-wide identification of lineage and locus specific variation associated with pneumococcal carriage duration. eLife 6, e26255 (2017).

59. Shaw, J. & Yu, Y. W. Rapid species-level metagenome profiling and containment estimation with sylph. Nat. Biotechnol. 43, 1348–1359 (2025).

60. Langmead, B. & Salzberg, S. L. Fast gapped-read alignment with Bowtie 2. Nat. Methods 9, 357–359 (2012).

61. Danecek, P. et al. Twelve years of SAMtools and BCFtools. GigaScience 10, giab008 (2021).

62. Page, A. J. et al. SNP-sites: rapid efficient extraction of SNPs from multi-FASTA alignments. *Microb*. Genomics 2, e000056 (2016).

63. Hunt, M. et al. AllTheBacteria - all bacterial genomes assembled, available and searchable. 2024.03.08.584059 Preprint at 10.1101/2024.03.08.584059 (2024).

64. Mikheenko, A., Prjibelski, A., Saveliev, V., Antipov, D. & Gurevich, A. Versatile genome assembly evaluation with QUAST-LG. Bioinformatics 34, i142–i150 (2018).

65. Schwengers, O. et al. Bakta: rapid and standardized annotation of bacterial genomes via alignment-free sequence identification: Find out more about Bakta, the motivation, challenges and applications, here. *Microb*. Genomics 7, 000685 (2021).

66. Schwengers, O. Bakta database. Zenodo 10.5281/ZENODO.4247252 (2025).

67. Tonkin-Hill, G. et al. Producing polished prokaryotic pangenomes with the Panaroo pipeline. Genome Biol. 21, 180 (2020).

68. Minh, B. Q. et al. IQ-TREE 2: New Models and Efficient Methods for Phylogenetic Inference in the Genomic Era. Mol. Biol. Evol. 37, 1530–1534 (2020).

69. Letunic, I. & Bork, P. Interactive Tree of Life (iTOL) v6: recent updates to the phylogenetic tree display and annotation tool. Nucleic Acids Res. 52, W78–W82 (2024).

70. Godoy, D. et al. Multilocus Sequence Typing and Evolutionary Relationships among the Causative Agents of Melioidosis and Glanders, Burkholderia pseudomallei and Burkholderia mallei. J. Clin. Microbiol. 41, 2068–2079 (2003).

71. Currie, B. J., Fisher, D. A., Anstey, N. M. & Jacups, S. P. Melioidosis: acute and chronic disease, relapse and re-activation. Trans. R. Soc. Trop. Med. Hyg. 94, 301–304 (2000).

72. Lees, J. A., Galardini, M., Bentley, S. D., Weiser, J. N. & Corander, J. pyseer: a comprehensive tool for microbial pangenome-wide association studies. Bioinformatics 34, 4310–4312 (2018).

73. Ooi, W. F. et al. The Condition-Dependent Transcriptional Landscape of Burkholderia pseudomallei. PLOS Genet. 9, e1003795 (2013).

74. Rongkard, P., et al. Dysregulated immunologic landscape of the early host response in melioidosis. JCI Insight 9, (2024).

75. Dobin, A. et al. STAR: ultrafast universal RNA-seq aligner. Bioinformatics 29, 15–21 (2013).

76. Liao, Y., Smyth, G. K. & Shi, W. featureCounts: an efficient general purpose program for assigning sequence reads to genomic features. Bioinformatics 30, 923–930 (2014).

77. Chen, Y., Chen, L., Lun, A. T. L., Baldoni, P. L. & Smyth, G. K. edgeR v4: powerful differential analysis of sequencing data with expanded functionality and improved support for small counts and larger datasets. Nucleic Acids Res. 53, gkaf018 (2025).

78. Zhang, Y., Parmigiani, G. & Johnson, W. E. ComBat-seq: batch effect adjustment for RNA- seq count data. NAR Genomics Bioinforma. 2, lqaa078 (2020).

79. Ritchie, M. E. et al. limma powers differential expression analyses for RNA-sequencing and microarray studies. Nucleic Acids Res. 43, e47 (2015).

80. Law, C. W., Chen, Y., Shi, W. & Smyth, G. K. voom: precision weights unlock linear model analysis tools for RNA-seq read counts. Genome Biol. 15, R29 (2014).

81. Badia-i-Mompel, P., et al. decoupleR: ensemble of computational methods to infer biological activities from omics data. *Bioinforma. Adv*. 2, vbac016 (2022).

82. Schubert, M. et al. Perturbation-response genes reveal signaling footprints in cancer gene expression. Nat. Commun. 9, 20 (2018).

83. Müller-Dott, S. et al. Expanding the coverage of regulons from high-confidence prior knowledge for accurate estimation of transcription factor activities. Nucleic Acids Res. 51, 10934–10949 (2023).

84. Grant, C. E., Bailey, T. L. & Noble, W. S. FIMO: scanning for occurrences of a given motif. Bioinformatics 27, 1017–1018 (2011).

85. Dudek, C.-A. & Jahn, D. PRODORIC: state-of-the-art database of prokaryotic gene regulation. Nucleic Acids Res. 50, D295–D302 (2022).

