## Supplementary information for "Integrating bacterial genomics and host transcriptomics links *Burkholderia pseudomallei* variation to melioidosis mortality and neutrophil responses"

### Host-adjusted bacterial GWAS identifies pathogen determinants of melioidosis mortality and host immune response

**This Supplementary Information contains 8 Supplementary Figures and 7 Supplementary Tables.**

**Supplementary Figure 1: Robustness of the lineage-mortality association across all combinations of clinical covariates**

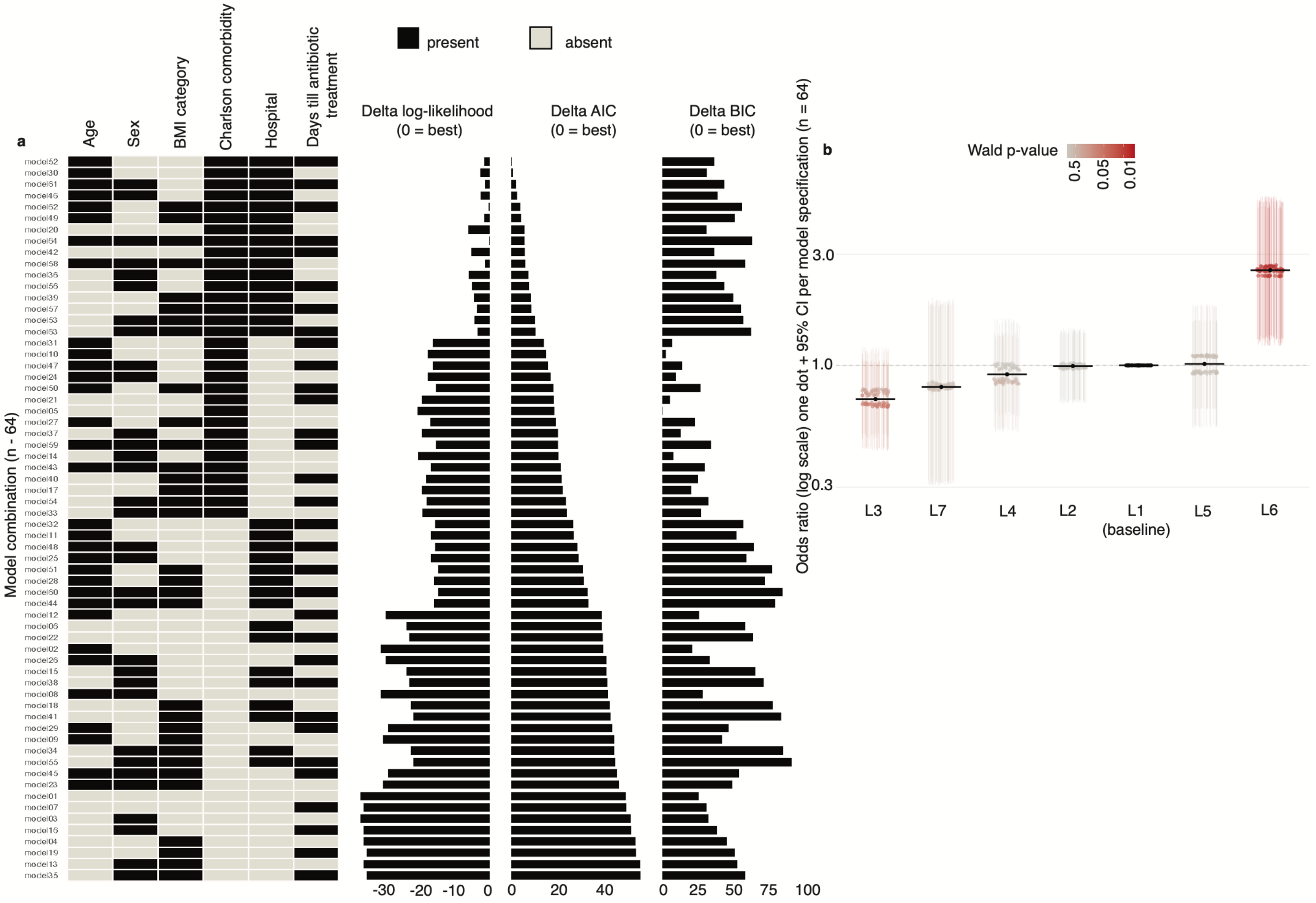


**(a)** Specification analysis of the 64 possible logistic regression models of 28-day mortality, each including infecting lineage plus a different combination of six clinical covariates: age, sex, BMI, Charlson comorbidity index, hospital, and days from admission to appropriate antibiotic treatment. **Left**: model specification (model 1-64) with filled cells indicate covariate included in that model (black, present; grey, absent). **Right**: model fit for each specification, expressed as the difference in log-likelihood, Akaike information criteria (ΔAIC), and in Bayesian information criteria (ΔBIC) relative to the best-fitting model. **(b)** Odds ratios for 28-day mortality by infecting lineage across all 64 model specifications, with lineage 1 as the baseline. Each vertical line with a dot is the odds ratio and 95% confidence interval from one specification, coloured by Wald p-value. Dashed horizontal line marks an odds ratio of 1: no difference from lineage 1. The elevated mortality with lineage 6 was consistent across every covariate combination tested, whereas the remaining lineages did not differ significantly from lineage 1 in any specification.

#### Supplementary Figure 2: Association between bacterial lineage and mortality in the Cox proportional hazards model

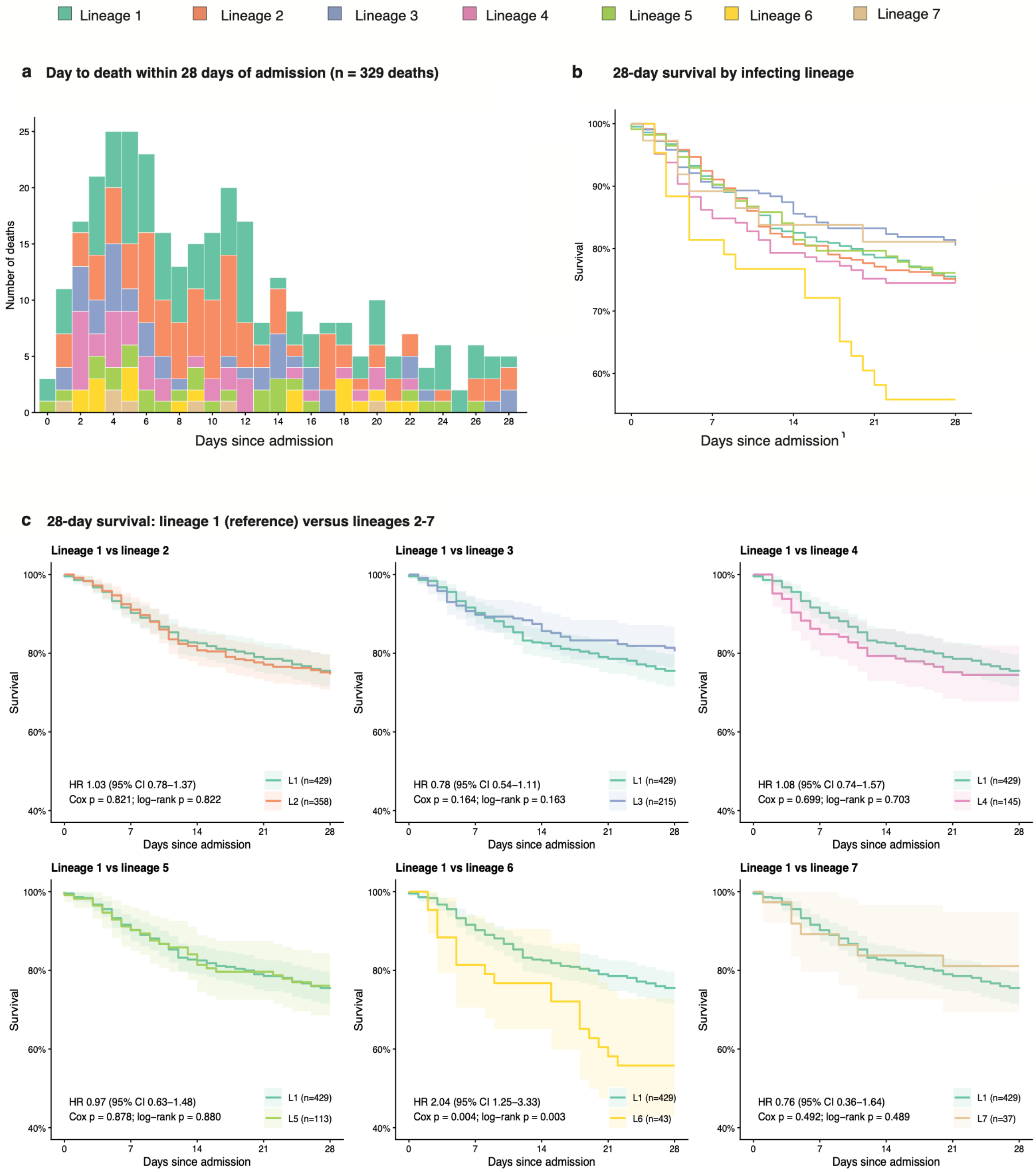


**(a)** Distribution of time from hospital admission to death among patients who died within 28 days (n = 329), in 1-day bins, with bars coloured by the lineage of the infecting isolate. Deaths were concentrated in the first two weeks, peaking at days 4-6. **(b)** Kaplan-Meier estimates of 28-day survival for all patients, stratified by infecting lineage. **(c)** Pairwise comparison of 28-day survival between lineage 1 (reference, teal) and each of lineages 2-7. Solid lines show Kaplan-Meier survival estimates and shaded bands at 95% CI.

**Supplementary Figure 3: Quantile-quantile plots of genome-wide association test statistics under three models**

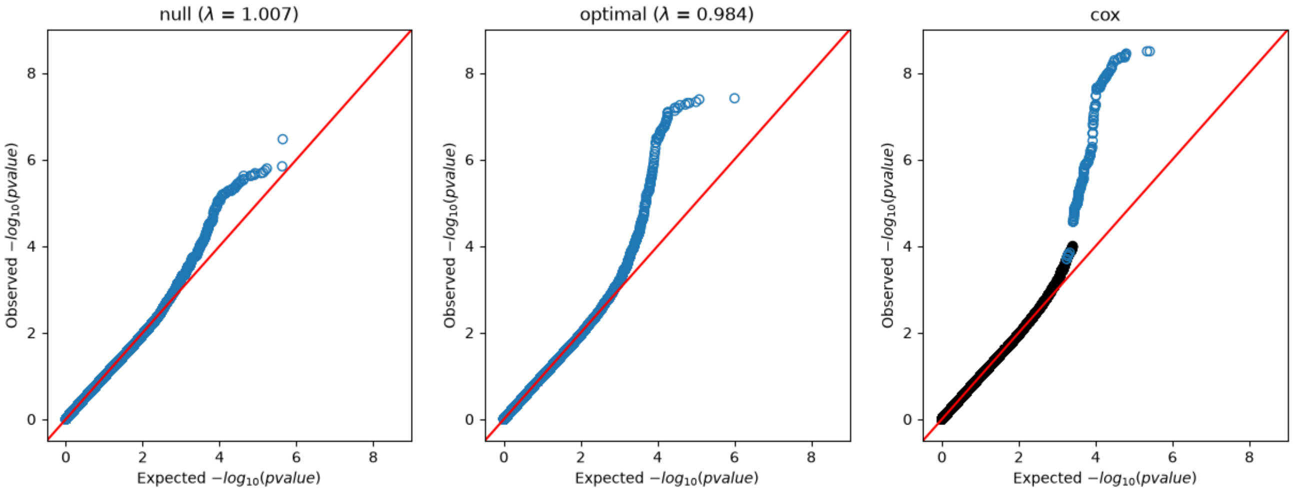


Quantile-quantile plots concern three models: (**left**) the LMM null model, adjusted for bacterial lineage only, (**middle**) the LMM optimal model, additionally adjusted for host age, sex, Charlson Comorditiy Index, and BMI category, and (**right**) the Cox GWAS using time to death as a continuous outcome variable censored at 28 days, adjusted for the same clinical covariates as the optimal model. Plots show observed versus expected $-\log_{10}(p)$ for all unitigs tested as blue circles; as only a subset of all unitigs were tested in Cox GWAS, these were shown as blue circles and black dots show the expected and observed values of untested unitigs in the LMM optimal model on which Cox GWAS was based. Red diagonals indicate the expectation under the null hypothesis of no association. Adjusting for lineage alone was sufficient to control inflation (genomic inflation factor $\lambda = 1.007$), and adding host sex, age, BMI category and Charlson comorbidity index left the bulk of the distribution well calibrated ($\lambda = 0.984$), with departure from expectation confined to the extreme upper tail – consistent with the genuine association signals rather than residual confounding by bacterial population structure. The Cox model was only run on unitigs pre-selected for $p < 10^{-4}$ in the LMM optimal model, to reduce computation load, so its departure from the diagonal is therefore expected by ascertainment and does not indicate inflation.

**Supplementary Figure 4 Distribution of mortality-associated unitigs and loci across bacterial phylogeny**

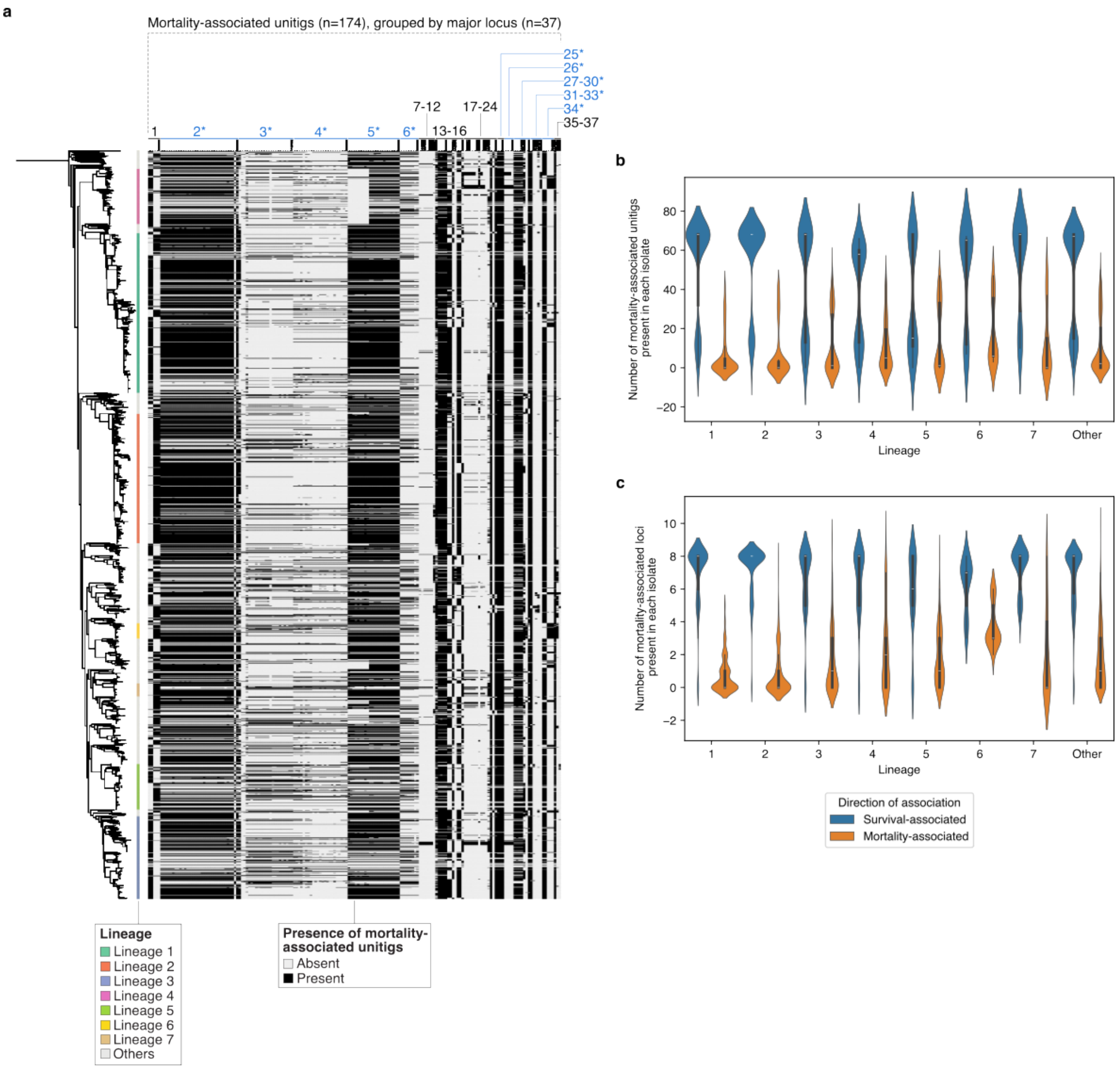


**(a)** Accompanying maximum-likelihood phylogeny of the 2,101 clinical *B. pseudomallei* isolates used in this study (*left*) are heatmaps showing isolate lineages (middle, coloured) and presence of mortality-associated unitigs ($p < 10^{-4}$; linear mixed model with adjustment for age, sex, BMI, and Charlson Comorbidity Index; heatmap on the right). Unitigs are grouped by major locus (locus numbers as labels at the top); indicated in blue are major loci whose unitigs all reach the $p < 10^{-5}$ suggestive significance threshold in Cox GWAS. **(b)** Violin plots showing the number of mortality-associated unitigs present in each isolate across the seven lineages, split between negative association (‘Survival-associated’) and positive association (‘Mortality-associated’). **(c)** Similar to (b), violin plots summarising the number of survival- and mortality-associated major loci present in each isolate across the seven lineages, with the presence of the unitig with the strongest association in Cox GWAS being a proxy for major locus presence.

**Supplementary Figure 5 Sensitivity analysis of bacterial unitigs associated with mortality.**

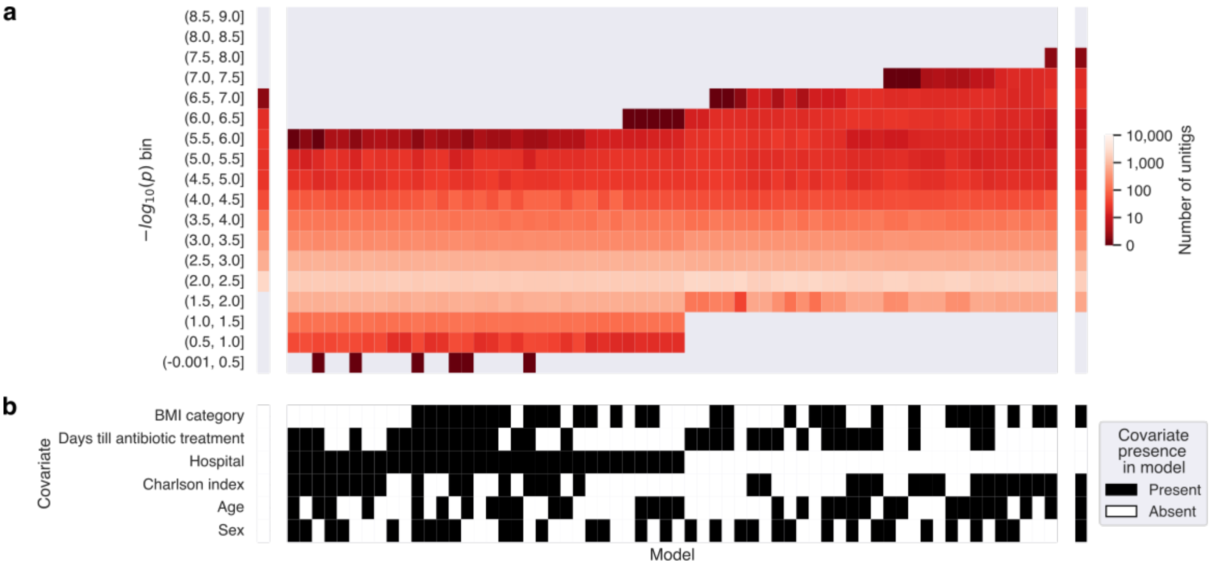


(**a**) Heatmap showing how genome-wide association results changed as the set of host covariates changes across models; all analyses used linear mixed models, fitted bacterial lineage as a random effect, and used a set of 1,821 patients for which all covariate values were defined. The heatmap shows the significance levels of unitigs with $p < 0.01$ in the null model (controlling for lineage only) and how p values change across models: each column is a model defined by a set of host covariates, each row is a bin of $-\log_{10}(p)$ values, and cells are shaded by the number of unitigs that fall in each bin (log colour scale, dark red to white: 0 to 10,000; grey bin indicate bins containing no unitigs). Apart from the null model (far left) and the optimal model (lineage plus sex, age, BMI category and Charlson comorbidity index; far right), models are ordered, left to right, by increasing maximum $-\log_{10}(p)$ attained in each model. (**b**) A key showing the covariate composition for each model, aligned column-wise with the heatmap. Filled (black) cells indicate the covariates included, among sex, age, Charlson Comorbidity Index, hospital, days from admission to antibiotic treatment, and BMI category.

**Supplementary Figure 6 Gene-level network of loci associated with melioidosis mortality**

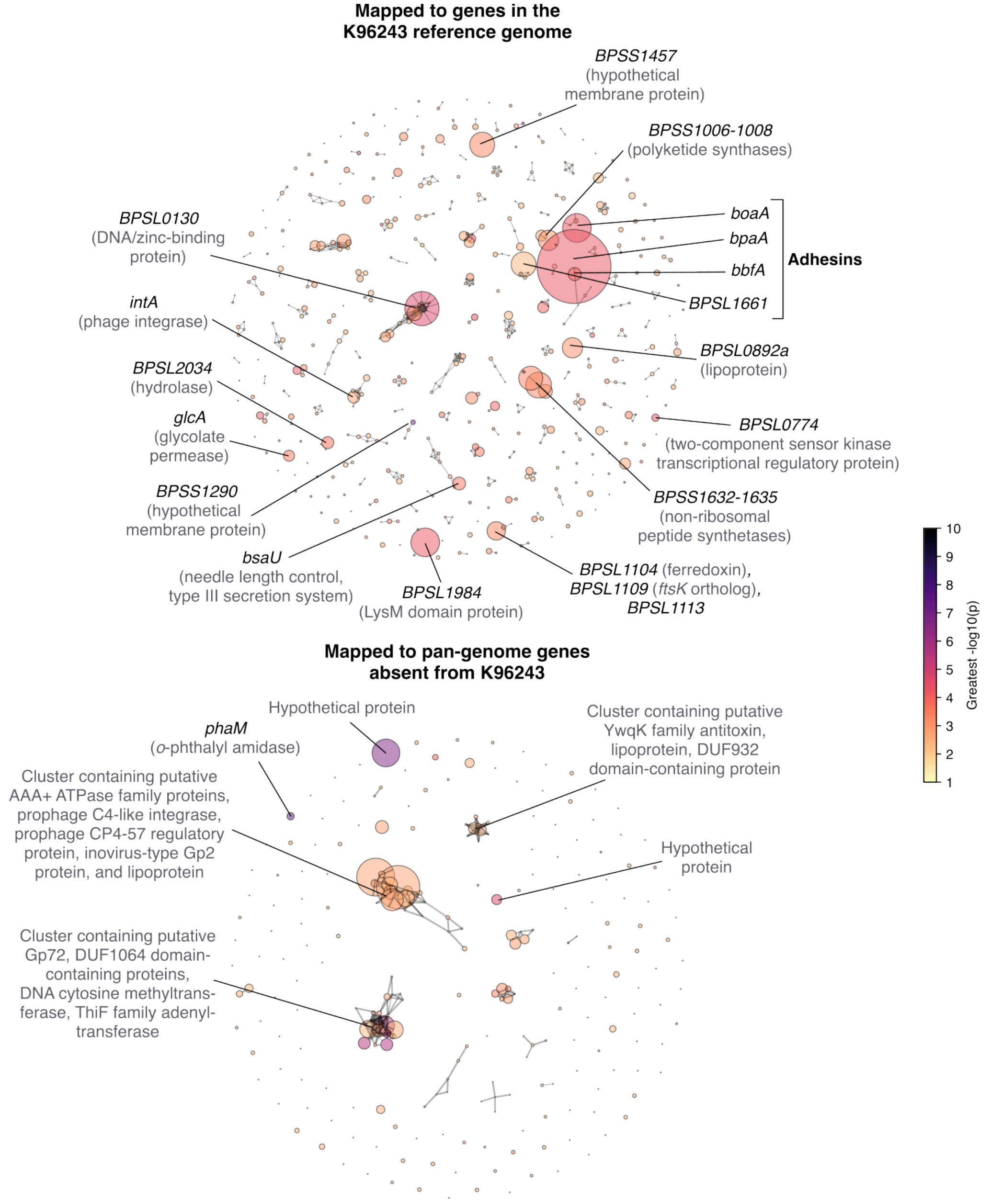


Network diagrams showing genes that unitigs reaching $p < 10^{-2}$ in the LMM optimal model map to – **top**: genes in K96243, **bottom**: pan-genome genes absent from K96243. Each node represents a gene, aggregating properties of unitigs mapped to it: node area is proportional to the number of mapped unitigs, node colour indicates the greatest $-\log_{10}(p)$ among the unitigs (colour bar, right). Edges connect genes within operon distance (7,000 bp); for the pan-genome diagram, the edges' thicknesses indicate the strength of evidence of such proximity calculated based on the proportion of assemblies possessing both joined genes within 7,000 bp. Key genes in the K96243 diagram are labelled with gene's name or locus tag, with the predicted product in parentheses if informative – the virulence genes *bpaA* (adhesin) and *bbfA* (biofilm formation protein) form the largest node pair. Key genes in the pangenome diagram are labelled with the genes' putative roles.

**Supplementary Figure 7 Peripheral immune cell composition in lineage 6 versus lineage 1 infection by cohort**

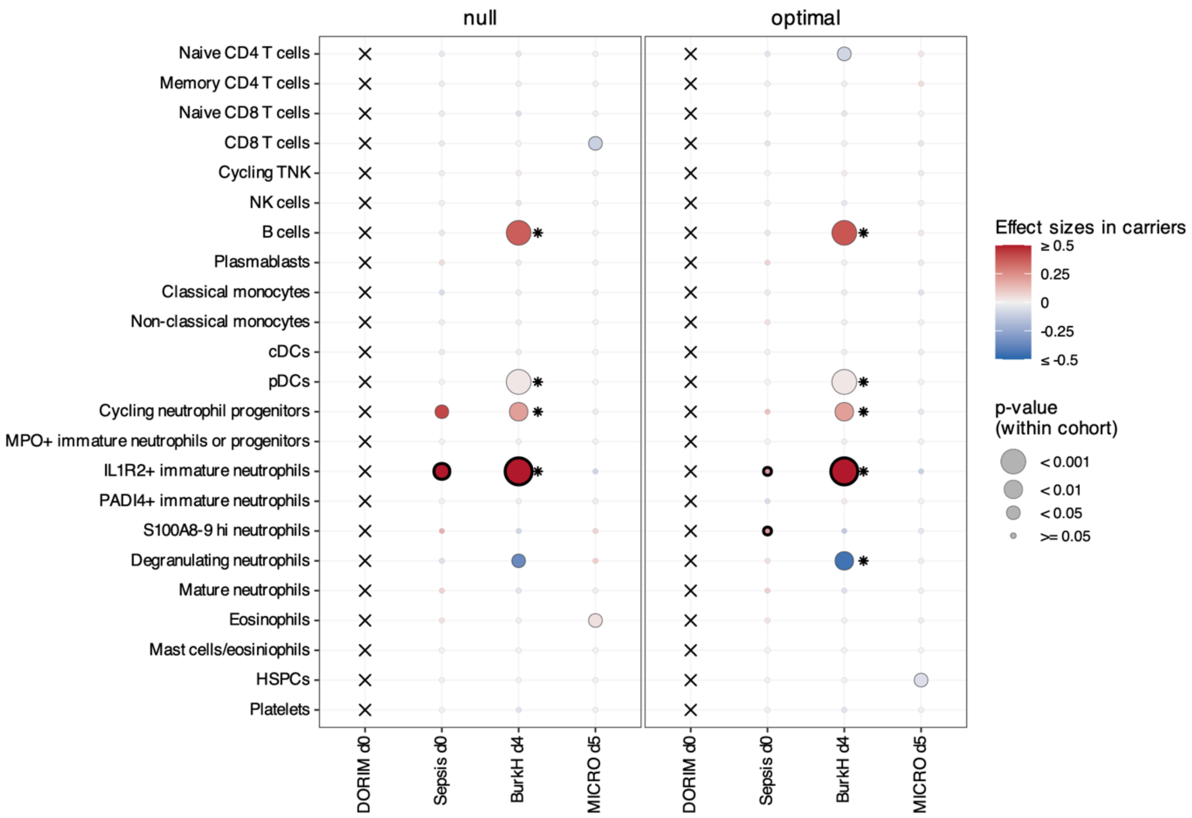


Differential abundance of 23 cell-type populations in patients infected with lineage 6 compared with lineage 1, estimated separately within each cohort. Rows are cell types; columns are cohorts with the sampling day indicated (DORIM, day 0; Ubon-Sepsis, day 0; BurkHostGEN, day 4; MICRO1501, day 5). **Left panel**: unadjusted models. **Right panel**: models adjusted for host covariates (sex, age, BMI category and Charlson comorbidity index). Point colour gives the regression coefficient for the lineage term from generalised linear models (red, higher abundance in lineage 6; blue, higher abundance in lineage 1), and point size gives the nominal p-value for that term. Crosses (x) denote cohort-cell type combinations for which no estimate could be obtained due to small (n < 5) sample sizes. BurkHostGEN yielded several associations that passed Benjamini-Hochberg correction (FDR = 0.05) – these are indicated with asterisks – however, these rest on only four lineage 6 patients and thus need cautious interpretation.

**Supplementary Figure 8 Genomic context of the mortality-associated ML34 locus**

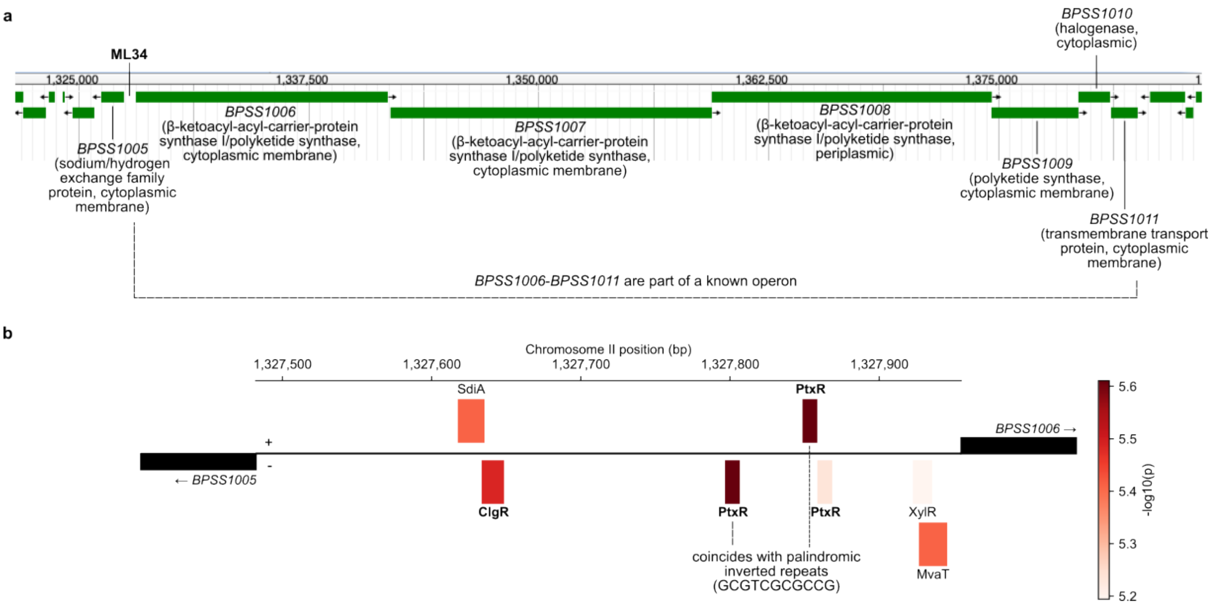


**(a)** Annotated gene map of the region (coordinates in bp on chromosome 2). Green bars are coding sequences, with arrowheads indicating the direction of transcription; predicted products and subcellular localisations are given in parentheses. BPSS1006-BPSS1011 form a known operon encoding a polyketide synthase cluster (dashed bracket). ML34 denotes the locus identified in the mortality association analysis. **(b)** Expanded view of the intergenic region between *BPSS1005* and *BPSS1006*, with putative transcription factor binding motifs on both strands predicted by FIMO using the PRODORIC database ($p < 10^{-5}$). *Pseudomonas aeruginosa* PtxR and *Corynebacterium glutaminum* ClgR binding motifs are highlighted as the motifs' respective transcription factors or nearest equivalents had > 40% identity with *Burkholderia pseudomallei* K96243 proteins – PtxR with *BPSS0166* LysR family transcriptional regulator (42.1%) and the *Corynebacterium glaucum* ClgR with *BPSS2142* MerR family transcriptional regulator (45.6%). The putative PtxR binding motifs coincided with palindromic inverted repeats GCGTCGCGCCG, one on each strand (1,327,859-1,327,849, plus strand; 1,327,796-1,327,806, minus strand). The same repeat pair occurs in the intergenic region between *BPSS0622* (enoyl-CoA hydratase) and *BPSS0623* (outer membrane efflux protein), although those genes are convergently transcribed, whereas *BPSS1005* and *BPSS1006* are divergent.

#### Supplementary Tables

##### Supplementary Table 1: Patient characteristics and host covariates

Clinical characteristics and host covariates for the 2,142 patients included in the study. Data are redacted or aggregated (**Table 1**) where necessary to protect participant confidentiality

##### Supplementary Table 2: Modified Charlson Comorbidity Index

Modified Charlson Comorbidity Index calculations based on the clinical variables recorded across the study cohorts included in this study.

| Clinical variable | Weight used in this study | Weight used by Quan et al. (2014) |
|---|---|---|
| Coronary heart disease | 0 | 0 |
| Congestive heart failure | 2 | 2 |
| Stroke | 0 | 0 |
| Chronic lung disease | 1 | 1 |
| Hepatitis (excluding moderate or severe liver disease) | 2 | 2 |
| Diabetes mellitus with presumed complications | 1 | 1 |
| Diabetes mellitus with no complications | 0 | 0 |
| Renal disease (eGPR < 60 mL/min) | 1 | 1 |
| Any malignancy | 2 | 2 |
| Peripheral vascular disease | N/A | 0 |
| Dementia | N/A | 2 |
| Rheumatologic disease | N/A | 1 |
| Peptic ulcer disease | N/A | 0 |
| Hemiplegia or paraplegia | N/A | 2 |
| Moderate or severe liver disease | N/A | 4 |
| Metastatic solid tumour | N/A | 6 |
| HIV/AIDS | N/A | 4 |
| **Maximum aggregated score** | **9** | **24** |

##### Supplementary Table 3: Bacterial isolates and associated metadata

Bacterial isolates sequenced from study patients and their associated clinical and genomic metadata.

##### Supplementary Table 4: Quality control of whole-genome sequenced bacterial isolates

Summary of quality-control metrics for the whole-genome sequenced *B. pseudomallei* isolates included in the study.

**Supplementary Table 5: Pairwise SNP distances within and between bacterial lineages**
Summary of pairwise SNP distances within each bacterial lineage and across the *B. pseudomallei* population

Whitney U test per lineage versus all

| Population/Lineage | N pairs | Median (Q1 – Q3) | BH adjusted p value |
|---|---|---|---|
| Population wide | 3,389,106 | 17,040 (15,559 - 18,210) | N/A |
| L 1 | 99,681 | 2,098 (779 - 2,860) | $<2 \times 10^{-16}$ |
| L 2 | 65,703 | 204 (151 - 316) | $<2 \times 10^{-16}$ |
| L 3 | 26,796 | 3,269 (2366 - 3,990) | $<2 \times 10^{-16}$ |
| L 4 | 11,935 | 3,902 (245 – 5,045) | $<2 \times 10^{-16}$ |
| L 5 | 8,128 | 4,610 (158 - 5,638) | $<2 \times 10^{-16}$ |
| L 6 | 903 | 102 (69 - 392) | $<2 \times 10^{-16}$ |
| L 7 | 666 | 63 (34 - 79) | $<2 \times 10^{-16}$ |

**Supplementary Table 6: Recombination and mutation estimates across bacterial lineages**
Summary of recombination relative to mutations (r/m) detected within each bacterial lineage.

Post-hoc pairwise Wlicoxon rank-sum test (BH-adjusted p-values)

| group 1 | group 2 | group1 median r/m (Q1-Q3) | group 2 median r/m (Q1-Q3) | BH adjusted p value |
|---|---|---|---|---|
| L 2 | L 1 | 0.428571 | 0.142857 | $1.60 \times 10^{-5}$ |
| L 4 | L 1 | 0.777778 | 0.142857 | $1.60 \times 10^{-5}$ |
| L 5 | L 4 | 0.130435 | 0.777778 | $1.60 \times 10^{-5}$ |
| L 5 | L 2 | 0.130435 | 0.428571 | $3.06 \times 10^{-5}$ |
| L 4 | L 3 | 0.777778 | 0.3 | $1.47 \times 10^{-4}$ |
| L 3 | L 2 | 0.3 | 0.428571 | $2.26 \times 10^{-3}$ |
| L 7 | L 4 | 0.2 | 0.777778 | $2.2 \times 10^{-3}$ |
| L 7 | L 2 | 0.2 | 0.428571 | $1.14 \times 10^{-2}$ |
| L 5 | L 3 | 0.130435 | 0.3 | $5.30 \times 10^{-2}$ |
| L 3 | L 1 | 0.3 | 0.142857 | $1.78 \times 10^{-1}$ |
| L 7 | L 3 | 0.2 | 0.3 | $1.78 \times 10^{-1}$ |
| L 6 | L 4 | 0.269231 | 0.777778 | $1.78 \times 10^{-1}$ |
| L 6 | L 5 | 0.269231 | 0.130435 | $1.78 \times 10^{-1}$ |
| L 7 | L 6 | 0.2 | 0.269231 | $2.91 \times 10^{-1}$ |
| L 4 | L 2 | 0.777778 | 0.428571 | $3.31 \times 10^{-1}$ |
| L 5 | L 1 | 0.130435 | 0.142857 | $3.36 \times 10^{-1}$ |
| L 6 | L 2 | 0.269231 | 0.428571 | $3.36 \times 10^{-1}$ |
| L 6 | L 1 | 0.269231 | 0.142857 | $3.61 \times 10^{-1}$ |
| L 7 | L 1 | 0.2 | 0.142857 | $6.67 \times 10^{-1}$ |
| L 6 | L 3 | 0.269231 | 0.3 | $7.07 \times 10^{-1}$ |
| L 7 | L 5 | 0.2 | 0.130435 | $9.17 \times 10^{-1}$ |

**Supplementary Table 7 Mortality-associated bacterial GWAS hits across host-adjusted models.**
Summary of GWAS-significant unitigs and loci identified across models, including the corresponding host covariates and the gain in statistical power associated with host adjustment.
